# Methylome-wide association studies of traumatic injury identifies differential DNA methylation of synaptic plasticity and GABAergic-signalling

**DOI:** 10.1101/2023.11.13.566835

**Authors:** Jodie P. Brown, Sally Marshall, Rosie M. Walker, Archie Campbell, Caroline Hayward, Andrew M. McIntosh, Kathryn L. Evans, Pippa A. Thomson

**Affiliations:** Edinburgh Medical School, University of Edinburgh; Centre for Genomic and Experimental Medicine, Institute of Genetics and Cancer, University of Edinburgh; School of Psychology, University of Exeter, Exeter, United Kingdom; MRC Human Genetics Unit, Institute of Genetics and Cancer, University of Edinburgh; Centre for Clinical Brain Sciences (Division of Psychiatry), University of Edinburgh

## Abstract

Traumatic physical injury is often associated with psychological trauma and is a risk factor for major depressive disorder (MDD). In Generation Scotland traumatic injury was significantly associated with recurrent major depression (OR = 2.10, 95% CI 1.33-3.33, *P*LJ=LJ0.0016). and schizotypal symptoms, particularly disorganised thought (βLJ=LJ0.111, 95% CI 0.049-0.177, *P*LJ=LJ0.0004). We performed methylome-wide analyses of traumatic injury in individuals with MDD and controls separately to investigate the link between traumatic injury and MDD. Nominally significant differences in differential DNA methylation between MDD and control groups were identified at 40 003 CpG sites (p < 0.05). Individuals with recurrent MDD showed significantly higher levels of DNA methylation associated with traumatic injury at CpG sites at the first exon and lower levels at exon boundaries, this was significant different to the association pattern at these sites in controls (mean difference in M-value = 0.0083, *P* = 21.1×10^-10^, and -0.0125, *P* = 2.1×10^-174^, respectively). Analyses at the level of CpG site, genes and gene ontologies implicated dysregulation of processes related to synaptic plasticity, including dendrite development, excitatory synapse and GABAergic signalling (normalised enrichment values > 2, FDR q-values < 0.01). Enrichment analyses for regional brain-expression in the recurrent MDD group highlight the limbic lobe and supraoptic nuclei (recurrent MDD FWER = 0.028 and 0.034, respectively). These results suggest that traumatic injury is associated with patterns of DNA methylation differentially in individuals with MDD compared to controls, highlighting the need for novel analysis approaches.

## Introduction

Major depressive disorder (MDD) is a multifactorial disorder with highly polygenic heritability and strong environmental risk factors including stressful life events and trauma. There is evidence that genetic predisposition and adverse life events interact to determine individual risk. Differential DNA methylation is associated with both self-reported traumatic events and MDD.^1, 2^ This suggests that DNA methylation may be used to identify the biological pathways leading to increased risk of MDD.

Studying environmental factors can be problematic, as they could be a cause or consequence of MDD. To counteract this issue, some have grouped life events into dependent/independent ^3-10^, but relatively few have looked objectively at external traumatic risk factors, such as traumatic injury. These can be measured objectively, in terms of the injury sustained, but can also reflect an individual’s perception of the seriousness of the incident. The perception of trauma may be important for poor mental health outcomes and treatment response.^11, 12^

Traumatic injury is a physical injury that requires urgent medical attention and can be the result of road traffic accidents, falls, violence, or sports injuries. According to the WHO, traumatic injury affects tens of millions of individuals globally each year resulting in significant long-term disability.^13^ Approximately 30-40% of survivors of traumatic injury develop increasing psychological problems including 9%-15% of whom develop MDD.^12, 14-16^ A prior history of mental health issues, persistent rumination about the trauma, perceived threat to life, lack of social support and persistent physical problems are predictors of poor mental health outcomes.^16-19^ Increased understanding of the effects of physical trauma on mental health has led to the development of a number of screening tools for the prediction of individuals at increased risk of generalised anxiety disorder, depression or post-traumatic stress disorder (PTSD)^20-23^, and to the development of NHS guidance for psychological support after trauma even in the absence of physical injury. Response to treatment of psychological distress is, however, heterogeneous.^15, 24^

We hypothesised that: i) self-reported traumatic injury may be associated with differential methylation and may identify biological processes that may mediate the effect of trauma, ii) investigating differential methylation in both individuals with diagnoses of lifetime major depression and healthy controls separately may identify the individual differences in response to trauma leading to mental health problems.

A traumatic injury phenotype incorporating both physical injury (bone fracture) and self-reported psychological distress was derived in the Generation Scotland cohort.^25, 26^ Preliminary phenotypic analyses confirmed a significant association between a positive traumatic injury response and lifetime diagnoses of MDD. Methylome-wide association studies (MWAS) were performed for differential DNA methylation associated with traumatic injury in three groups: individuals without psychiatric diagnoses (Control), combined single or recurrent episode MDD (MDD) and recurrent only MDD (rMDD). The results of the MWAS were analysed to investigate pathway-specific enrichment and to identify differences in differential DNA methylation between the groups (methylome-wide x environment interaction studies, MWEIS: xMDD, xrMDD) and associated with traumatic injury across all individuals (META). Pathway-specific polygenic risk scores were used to assess the association of traumatic injury-related biological processes with MDD in an independent subgroup of Generation Scotland participants. Finally, regional-brain expression patterns over-represented in the genes associated with differential DNA methylation were investigated.

## Material and methods

### Study cohort

Generation Scotland is a population-based cohort of approximately 24 000 participants.^25-27^ Participants were recruited via general medical practices in Scotland between 2006 and 2011 and completed physical test and health questionnaires (data available on request: http://www.generationscotland.co.uk). All participants have given informed consent. Ethics approval for Generation Scotland was given by the NHS Tayside committee on research ethics (reference 05/S1401/89).

### Phenotypes

Information on traumatic incident that resulted in bone fracture was collected as part of the Generation Scotland questionnaire on musculoskeletal history in which participants were asked whether they had ever broken a bone and, if so, whether the fracture had occurred “in a road traffic accident or other serious incident”. In total, approximately 11 800 participants answered whether they had broken a bone, of whom 11 387 reported at least one fracture. An individual was classified as having had traumatic injury if they responded yes to the follow-up question (N=1 010).

A diagnosis of major depressive disorder (MDD) was made using the structured clinical interview for DSM-IV disorders (SCID).^28^ Participants who answered yes to either of the two screening questions were invited to continue the interview, which provided information on the presence or absence of a lifetime history of MDD and number of depressive episodes. Participants who answered no to both screening questions or who completed the SCID but did not meet the criteria for depression were assigned control status. Participants with diagnoses of schizophrenia or bipolar disorder were excluded from the analyses.

The age and sex of participants, and additional questionnaires to assess cognitive ^29-31^ ability and personality ^32^ and traits reflecting symptoms associated with depression (GHQ)^33^, bipolar disorder (MDQ)^34^ and schizophrenia (SPQ-BR) were collected at baseline.^35, 36^

### Phenotypic analyses

Logistic regression analyses of traumatic injury were performed in R (^37^) using the *glm* function in MASS.^38^ The phenotypic sample consisted of 2 946 individuals including 398 MDD individuals (72% female) and 2 548 controls (59% female). Phenotypic analyses were adjusted for sex and age at baseline. Variables from SPQ, MDQ and GHQ were log2 transformed to improve the normal distribution of the residual variance. Coefficients are reported on the non-log2 scale.

### MWAS & MWEIS

Whole blood DNA methylation was profiled using the Infinium MethylationEPIC BeadChip (Illumina Inc.) in two sets of GS participants at two separate times by the Wellcome Trust Clinical Research Facility, Edinburgh. The natural discovery sample comprised 5 190 individuals and the natural replication sample included 4 583 individuals, as described previously.^1, 39, 40^ The two samples were normalised, and the data was converted to M-values. Individuals in the replication sample were unrelated to those in the discovery set (SNP-based relatedness < 0.05). The data was quality controlled prior to the analyses to remove poor-performing probes, sex chromosome probes, individuals with unreliable self-report data, suspected XXY genotype, as described in Walker et al.^41^

Individual MWAS were performed for traumatic injury in three groups: controls, MDD and a subset of MDD with evidence for recurrent episodes (rMDD, see Supplementary Table 1). *Limma* was used to calculate empirical Bayes moderated t-statistics from which the *P*-values were obtained.^42^ Covariates were fitted to the discovery and replication sample following Walker et al.^41^ The discovery sample M-values were pre-corrected for relatedness, estimated cell count proportions (granulocytes, natural killer cells, B lymphocytes, CD4+ T lymphocytes, and CD8+ T lymphocytes) and processing batch. This sample was also corrected for age, sex, smoking status, pack years, 20 methylation PCs and 20 genetic PCs. The replication sample M-values were corrected for age, sex, smoking status, pack years, estimated cell count proportions, processing batch, 20 methylation principal components and 20 genetic principal components.

The MWAS results for controls and MDD were further meta-analysed using an inverse standard error-weighted fixed model implemented in METAL to give the data set META. The meta-analysis included 997 608 loci from 4 308 individuals.

MWEIS were performed using a two-sample Z-test to test for significant differences in the effect size (fold-change in the ratio of unmethylated to methylated CpG sites) of the association with traumatic injury in the healthy control versus MDD groups to give the datasets xMDD and xrMDD.

A genome-wide significant *P* threshold, *P*LJ<LJ9.42 x 10^-8^, was applied to all MWAS and MWEIS results within the study, following Mansell et al.^43^.

### Functional enrichment for *MATN2* and *ZEB2* co-expression and protein-interaction network analyses

Co-expression partners were identified using GeneCards.^44^.The top 25 human protein-interaction network partners were identified using STRING.^45^ Enrichment analyses were performed in STRING using the whole genome as the background. STRING assesses enrichment within categories defined from external sources and *P*-values are corrected for multiple testing within each category using the Benjamini–Hochberg procedure. The top five most false discovery rate (FDR)-significant functional enrichments are reported.

### Gene set enrichment analyses

CpG sites were mapped to the nearest genes based on nearest transcriptional start site within 10 Kb. Over-representation analyses for gene ontology terms were performed using *WebGestaltR*.^46^ Gene ontologies were considered significant at an FDR q < 0.05 and a minimum of five genes in the target list.

Gene set expression enrichment in adult human brain regions was assessed by hypergeometric tests using *ABAEnrichment*^47^ at three MWAS *P* -value thresholds (T1 p<2.668×10^-6^, -log*P* = 5.5738; T2 *P* < 1×10^-5^, -log*P* = 5; T3 *P* < 0.005, -log*P* = 2.3) and three expression quantiles: 0.5, 0.7, and 0.9. Brain regions were considered significant at a family-wise error rate (FWER) < 0.05.

### Polygenic risk scores (PRS) analyses

Polygenic risk scores were calculated from genome-wide summary statistics from a GWAS of adult trauma in UK Biobank (see Supplementary materials for methods) or the 2018 PGC MDD meta-analysis (excluding Generation Scotland, Wray et al.) using PRSice-2 (v2.3.5)^48^ PRSet functions to calculate genome-wide and pathway specific PRS. Independent SNPs were selected using clumping with a window of 250 kb, threshold *P* = 1 and an r^2^ = 0.1. PRS were derived for six *P* -value thresholds (0.00000005, 0.00001, 0.001, 0.05, 0.5, 1). Gene regions were defined using hg19 with 2 kb flanking regions 5’ and 3’ of the transcription start and stop sites. Significance was accepted at P < 0.05 using the competitive *P*-value derived after 10 000 set permutations. The competitive permutations test for enrichment of signals in the tested pathways compared to match numbers of post-clump SNPs selected from the whole genome (Base set). Analyses were corrected for age, sex and 10 genetic PCs.

PRS analyses were performed on two independent sets of unrelated individuals from Generation Scotland. The first set contained unrelated individuals who answered the traumatic injury question (SetA, N = 3 522, including: 3 185 controls and 327 individuals reporting traumatic injury; 3 010 controls and 512 MDD cases of which 253 were rMDD cases). These individuals contributed to the MWAS/MWEIS analyses. The second set of individuals had no information on traumatic injury (SetB, N = 1 515, including 1 274 MDD controls, 241 MDD cases of which 117 were rMDD cases). This second set were not included in the MWAS/MWEIS analyses and are unrelated to those individuals in the first set. Generation Scotland genome-wide genotype data were generated using the Illumina HumanOmniExpressExome-8 v1.0 DNA Analysis BeadChip (San Diego, CA, USA) and Infinium chemistry by the Clinical Research Facility, University of Edinburgh. Full details can be found in references^27, 49^

## Results

### Association between traumatic injury and psychiatric traits

Traumatic injury was significantly associated with a lifetime diagnosis of depression (*P* = 0.0048; OR = 1.67, 95% CI = 1.17-2.39) in those individuals with evidence of recurrent episodes (*P* = 0.0016; OR = 2.10, 95% CI = 1.33-3.33), but not in those reporting single episodes (*P* = 0.30; OR = 1.31, 95% CI = 0.78-2.20, Table 1). Traumatic injury was associated with symptom measures and personality traits relevant to aspects of psychiatric disease; with individuals reporting traumatic injury having significantly higher scores on the Mood disorder questionnaire and the Schizotypal Personality Questionnaire-Brief Revised scale, particularly for the disorganised factor (Table 1). Nominally significant association was also seen with the depression subscale of the General Health Questionnaire and the personality trait: digit symbol coding.^50, 51^

**Table 1.**
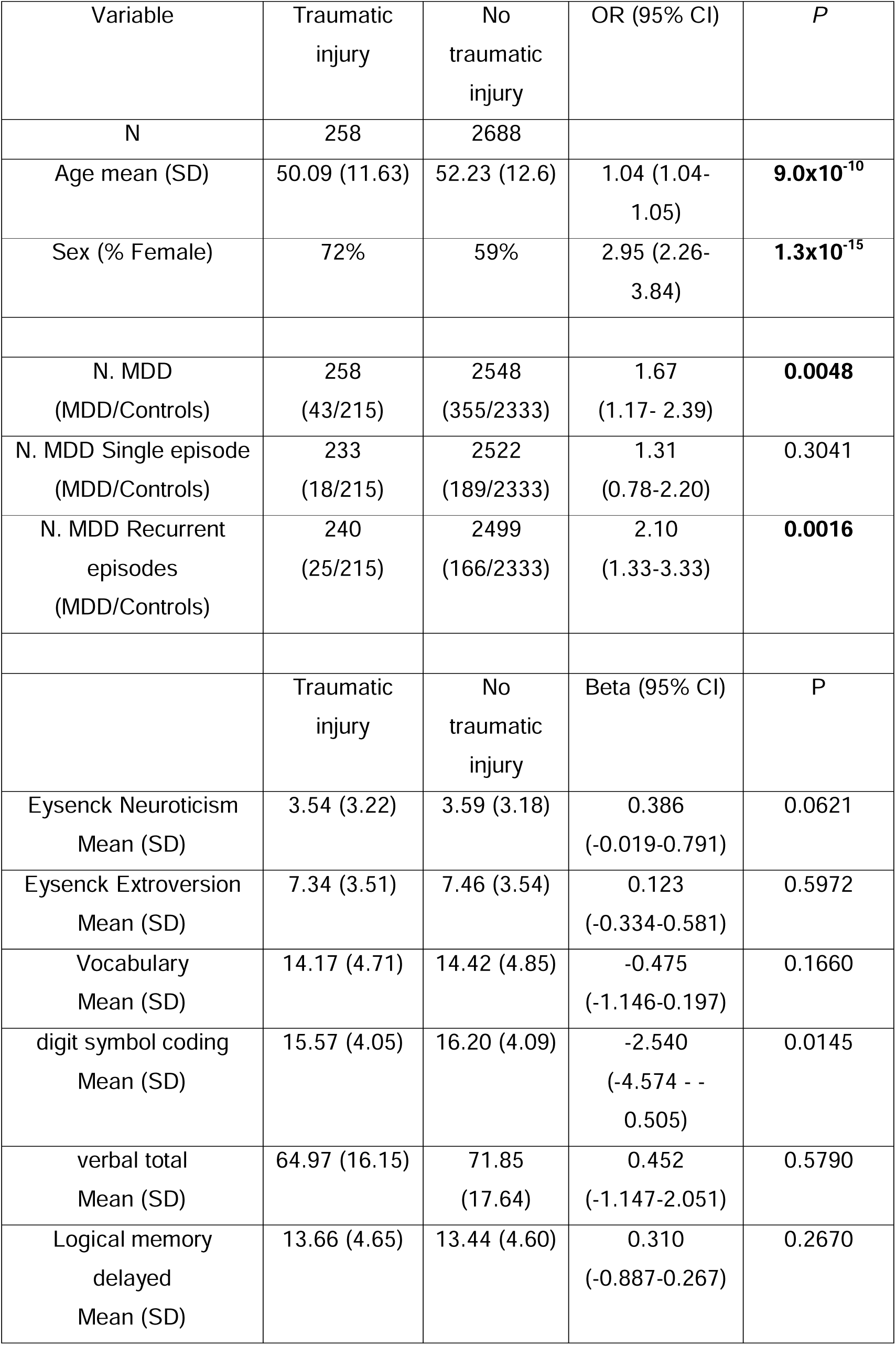
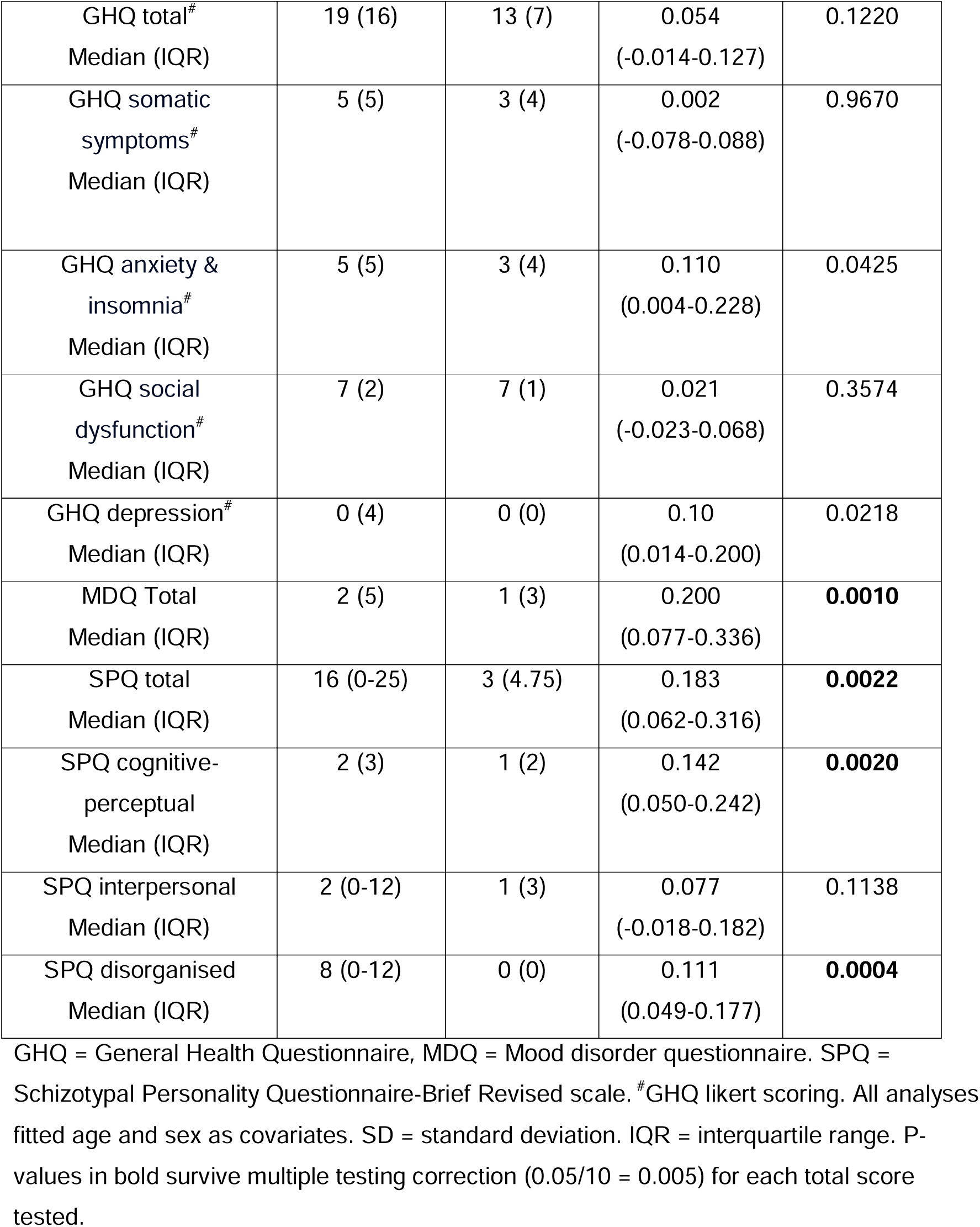
Phenotypic analyses of traumatic injury.

| Variable | Traumatic injury | No traumatic injury | OR (95% CI) | *P* |
| --- | --- | --- | --- | --- |
| N | 258 | 2688 |  |  |
| Age mean (SD) | 50.09 (11.63) | 52.23 (12.6) | 1.04 (1.04-1.05) | **9.0x10<sup>-10</sup>** |
| Sex (% Female) | 72% | 59% | 2.95 (2.26-3.84) | **1.3x10<sup>-15</sup>** |
| N. MDD (MDD/Controls) | 258 (43/215) | 2548 (355/2333) | 1.67 (1.17- 2.39) | **0.0048** |
| N. MDD Single episode (MDD/Controls) | 233 (18/215) | 2522 (189/2333) | 1.31 (0.78-2.20) | 0.3041 |
| N. MDD Recurrent episodes (MDD/Controls) | 240 (25/215) | 2499 (166/2333) | 2.10 (1.33-3.33) | **0.0016** |
|  | Traumatic injury | No traumatic injury | Beta (95% CI) | *P* |
| Eysenck Neuroticism Mean (SD) | 3.54 (3.22) | 3.59 (3.18) | 0.386 (-0.019-0.791) | 0.0621 |
| Eysenck Extroversion Mean (SD) | 7.34 (3.51) | 7.46 (3.54) | 0.123 (-0.334-0.581) | 0.5972 |
| Vocabulary Mean (SD) | 14.17 (4.71) | 14.42 (4.85) | -0.475 (-1.146-0.197) | 0.1660 |
| digit symbol coding Mean (SD) | 15.57 (4.05) | 16.20 (4.09) | -2.540 (-4.574 - -0.505) | 0.0145 |
| verbal total Mean (SD) | 64.97 (16.15) | 71.85 (17.64) | 0.452 (-1.147-2.051) | 0.5790 |
| Logical memory delayed Mean (SD) | 13.66 (4.65) | 13.44 (4.60) | 0.310 (-0.887-0.267) | 0.2670 |
| GHQ total<sup>#</sup><br>Median (IQR) | 19 (16) | 13 (7) | 0.054<br>(-0.014-0.127) | 0.1220 |
| GHQ somatic<br>symptoms<sup>#</sup><br>Median (IQR) | 5 (5) | 3 (4) | 0.002<br>(-0.078-0.088) | 0.9670 |
| GHQ anxiety &<br>insomnia<sup>#</sup><br>Median (IQR) | 5 (5) | 3 (4) | 0.110<br>(0.004-0.228) | 0.0425 |
| GHQ social<br>dysfunction<sup>#</sup><br>Median (IQR) | 7 (2) | 7 (1) | 0.021<br>(-0.023-0.068) | 0.3574 |
| GHQ depression<sup>#</sup><br>Median (IQR) | 0 (4) | 0 (0) | 0.10<br>(0.014-0.200) | 0.0218 |
| MDQ Total<br>Median (IQR) | 2 (5) | 1 (3) | 0.200<br>(0.077-0.336) | **0.0010** |
| SPQ total<br>Median (IQR) | 16 (0-25) | 3 (4.75) | 0.183<br>(0.062-0.316) | **0.0022** |
| SPQ cognitive-<br>perceptual<br>Median (IQR) | 2 (3) | 1 (2) | 0.142<br>(0.050-0.242) | **0.0020** |
| SPQ interpersonal<br>Median (IQR) | 2 (0-12) | 1 (3) | 0.077<br>(-0.018-0.182) | 0.1138 |
| SPQ disorganised<br>Median (IQR) | 8 (0-12) | 0 (0) | 0.111<br>(0.049-0.177) | **0.0004** |

GHQ = General Health Questionnaire, MDQ = Mood disorder questionnaire. SPQ = Schizotypal Personality Questionnaire-Brief Revised scale. <sup>#</sup>GHQ likert scoring. All analyses fitted age and sex as covariates. SD = standard deviation. IQR = interquartile range. P-values in bold survive multiple testing correction (0.05/10 = 0.005) for each total score tested.

### Genome-wide assessment of differential DNA methylation associated with traumatic injury

Genome-wide, 40,003 CpG sites, out of 756 971, showed nominally significant differences in the fold change of methylated to unmethylated CpG sites associated with traumatic injury in Control versus rMDD groups (xrMDD *P* < 0.05, 40 003/756, 971 = 0.052). Significant depletion of these sites was seen for CpG sites associated with the 1^st^ exon (3.2% v 24.4%, *P* < 1 x 10^-16^). Across all CpG sites there was no significant difference in the methylation levels in Control, rMDD or xrMDD (Table 2, Supplementary table 2). At sites associated with the first exon, however, the average DNA methylation was higher in individuals with rMDD who reported traumatic injury compared to those who did not report traumatic injury. This pattern was reversed in controls who reported traumatic injury compared to those not reporting traumatic injury (Table 2). The mirroring of the pattern was also seen at exon boundaries, with higher levels of DNA methylation in controls reporting versus not reporting traumatic injury and lower levels of DNA methylation in those with rMDD reporting versus not reporting traumatic injury.

**Table 2:** Mean difference in M-values at CpG sites with significant differences in the association of traumatic injury with differential methylation in the Control compared to rMDD groups.

| Feature | N. CpG sites<br>Genome | N $P < 0.05$<br>xrMDD | %<br>Genome/<br>xrMDD | Control<br>($\Delta M$-val) | rMDD<br>($\Delta M$-val) | xrMDD<br>($\Delta M$-val) | $P$<br>xrMDD |
| --- | --- | --- | --- | --- | --- | --- | --- |
| No annotation | 214550 | 11043 | 21.51 / 27.61 | -0.00068 | 0.00408 | **0.00477** | **$2 \times 10^{-4}$** |
| 1stExon | 243225 | 1292 | 24.38 / 3.23<sup>#</sup> | **-0.00051** | **0.00783** | **0.00835** | **$7 \times 10^{-62}$** |
| 3'UTR | 19499 | 996 | 1.95 / 2.49 | **-0.00151** | 0.00090 | **0.00241** | **$1 \times 10^{-10}$** |
| 5'UTR | 67698 | 3466 | 6.79 / 8.66 | -0.00013 | 0.00612 | **0.00625** | **$2 \times 10^{-18}$** |
| Body | 288428 | 14710 | 28.91 / 36.77 | -0.00054 | 0.00452 | **0.00506** | **$3 \times 10^{-3}$** |
| ExonBnd | 5310 | 255 | 0.53 / 0.64 | **0.00176** | **-0.01070** | -<br>**0.01246** | **$2 \times 10^{-174}$** |
| TSS1500 | 98047 | 5009 | 9.83 / 12.52 | -0.00026 | 0.00230 | **0.00256** | **$4 \times 10^{-3}$** |
| TSS200 | 60851 | 3232 | 6.10 / 8.08 | **-0.00070** | 0.00565 | **0.00635** | **$1 \times 10^{-16}$** |
| ALL | 997608 | 40003 |  | -0.00055 | 0.00392 | 0.00447 | **$7 \times 10^{-2}$** |

$\Delta M$ -val = difference in M-value. <sup>#</sup> Significantly depleted in xrMDD sites $P < 1 \times 10^{-16}$ ; Bold type = significantly different from 0 at $P < 0.05$ . Full details Supplementary table 2.

### Methylome-wide association and methylome-wide environment interaction studies of traumatic injury

The association of traumatic injury with DNA methylation was assessed separately in controls (Control, Supplementary Table 3) and in individuals with a lifetime diagnosis of MDD (MDD, Supplementary Table 4) and those with rMDD (Supplementary Table 5). Evidence of differential methylation in the control versus the MDD groups (MWEIS) was also investigated (xMDD and xrMDD, Supplementary Tables 6 & 7). Finally, meta-analysis of the Control and MDD groups was performed to identify consistent evidence for differential DNA methylation associated with traumatic injury across all individuals (META, Supplementary Table 8).

Two CpG sites showed methylome-wide significant differential DNA methylation in either the MWAS or MWEIS of traumatic injury (cg14764459, MWAS Control and META; cg02101279, MWEIS xrMDD; Supplementary Figure 1). Sixty-six CpG sites had p-values less than a suggestive significance level (*P* < 1 x 10^-5^, Supplementary Table 9). There were no shared CpG sites of suggestive significance between Control and either MDD group MWAS. However, three CpG sites show suggestive significance across multiple MDD MWAS/MWEIS groups (cg02101279, cg04640885, cg13395761). CpG site cg02101279, maps to a lincRNA *RP11-46E17.6*, and may show lower levels of DNA methylation in the rMDD individuals who reported traumatic injury. MWEIS analyses indicate that both the MDD and rMDD groups differ significantly from the Control group in the differential methylation levels associated with traumatic injury at this site, with higher levels of DNA methylation in the MDD groups and decreased DNA methylation in the control group reporting traumatic injury. The opposite pattern was observed for cg04640885, that maps to *ZEB2*, with higher levels of DNA methylation in the control group versus lower levels of DNA methylation in MDD, when they reported traumatic injury and in both MWEIS analyses (Figure 1a, 1c). Finally, cg13395761 maps within the *MATN2* and *RPL30* genes, and shows no evidence for differential DNA methylation in controls, but may show an association with higher levels of DNA methylation in the MDD individuals reporting traumatic injury (Figure 1b, 1c). Analysis of *MATN2* and *ZEB2* co-expression and protein-interaction networks suggests that they function in NOTCH signalling and negative regulation of transcription, respectively (Supplementary Tables 10-13).

**Figure 1:**
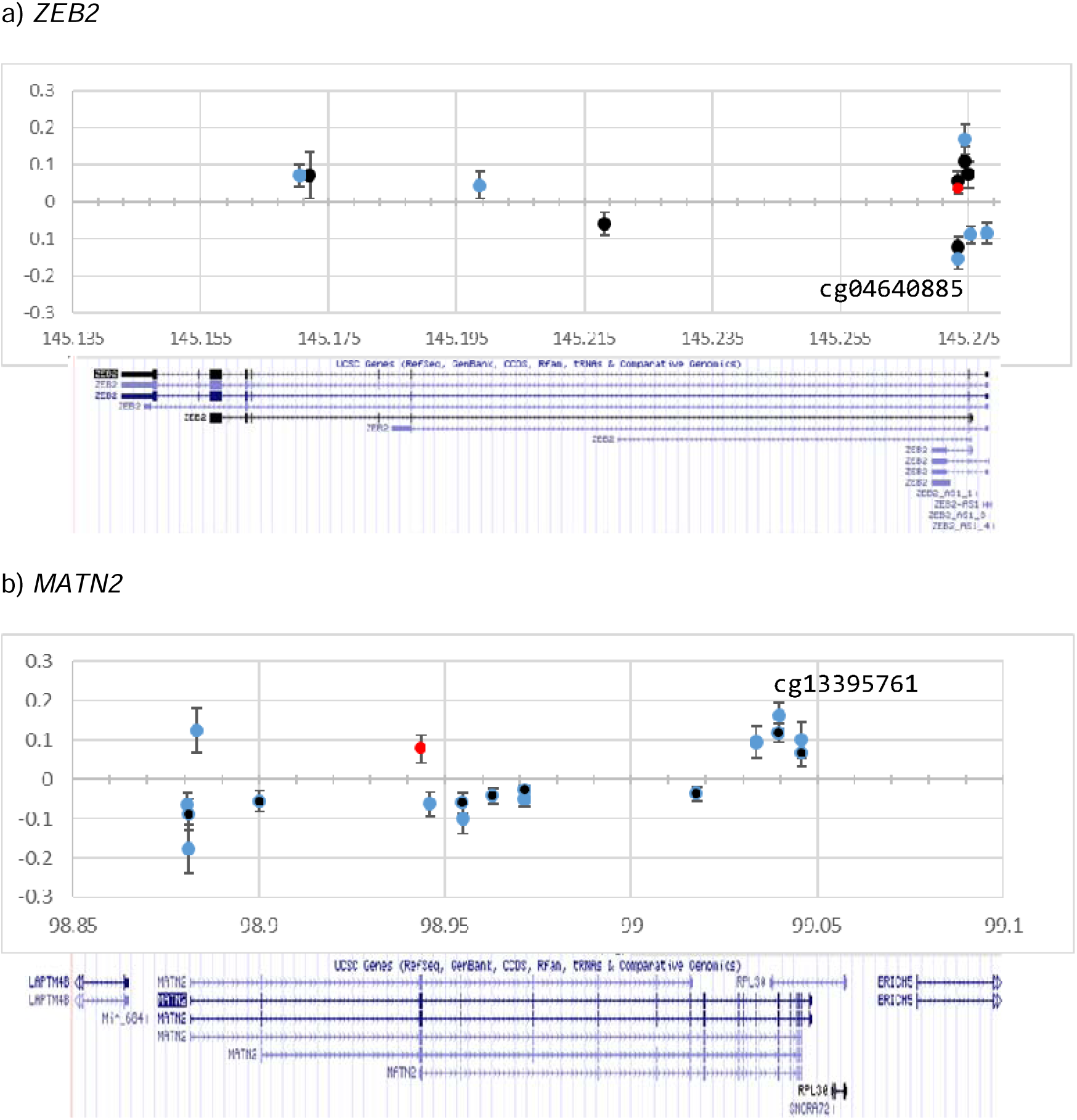
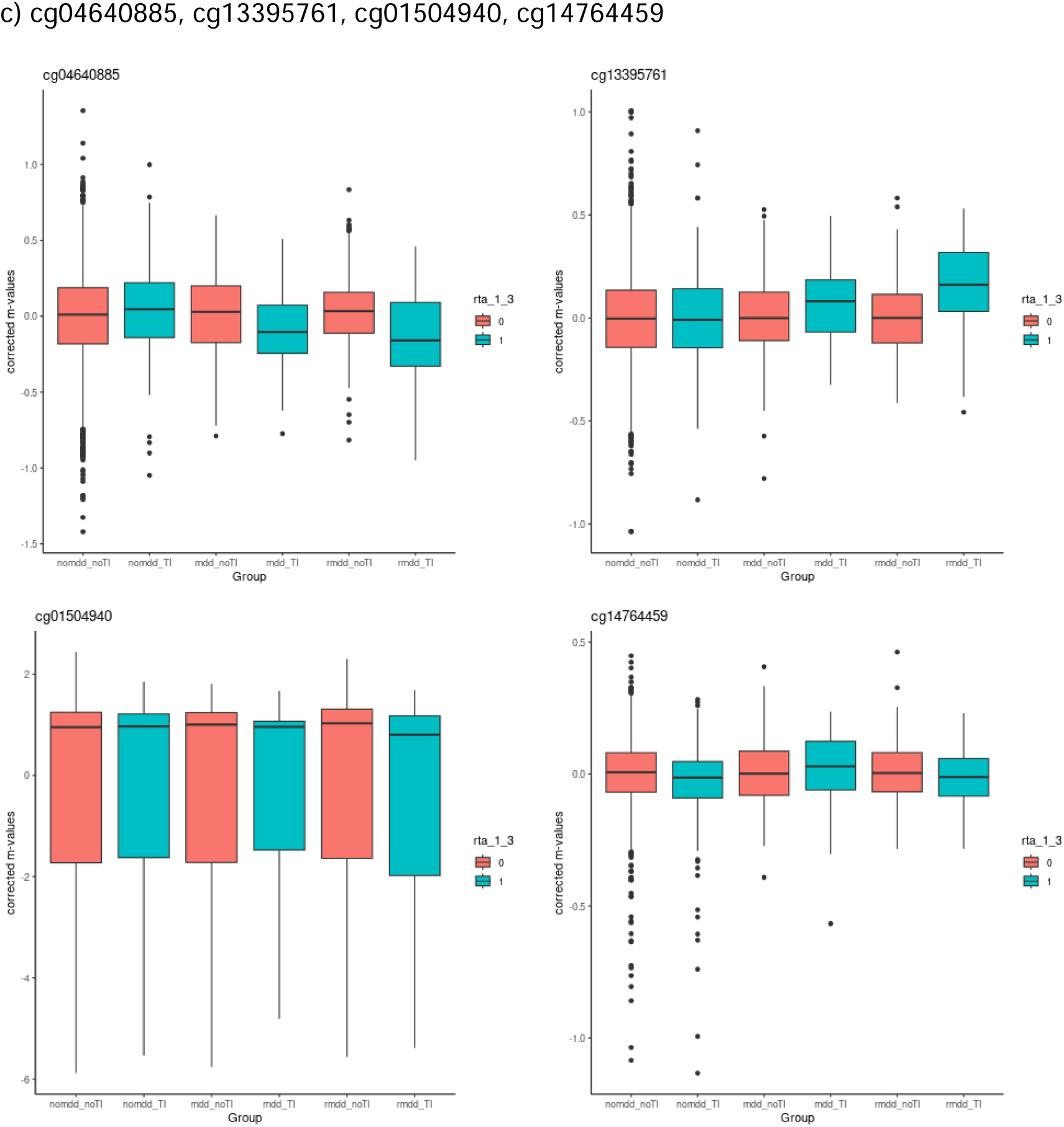
Difference in M-value at CpG sites with nominally significant differential methylation (*P* < 0.05) associated with traumatic injury in a) *ZEB2* and b) *MATN2*; for Control (red), any MDD (black) and rMDD (light blue). X-axis – position in Mbp. Y-axis – difference in M-value between individuals reporting traumatic injury and those not reporting traumatic injury, Error bars – standard error. Lower panel shows the genes in the region. c) Boxplot of M-values (corrected for covariates) for cg04640885 (*ZEB2*), cg13395761 (*MATN2*), cg01504940 (*MATN2*) and cg14764459 (*LOC100287944*), by group: Control_noTI, Control_TI, smdd_noTI, smdd_TI, rmdd_noTI, rmdd_TI.

These results are, however, not genome-wide significant and will require replication. There was little overlap between the genes mapping to CpG sites with suggestive significance in controls and rMDD (Supplementary Figure 2).

### MWAS/MWEIS gene set enrichment analyses

Gene set enrichment analyses, performed on the ranked lists of mapped genes in WebGestalt^46^, identified 808 gene ontology (GO) terms at an FDR significant q < 0.05 in at least one MWAS/MWEIS of traumatic injury, of which 205 terms were significant in all six (Control, MDD, rMDD, xMDD, xrMDD, META; Supplementary Table 14). Multicellular organismal response to stress (GO:0033555) was significant at an FDR q-value < 0.05 for all the analyses except xMDD, and showed the greatest enrichment of this term in the xrMDD analysis (33 leading edge genes, normalized enrichment values = 1.89, FDR q = 0.011).

Comparison of the gene ontology terms found to be enriched at FDR q < 0.05 across the MWAS and MWEIS shows that the majority are shared in the analyses of both Control and rMDD (325/563 GO terms for rMDD are shared with Control). However, the majority of these terms are also enriched in xrMDD, suggesting enrichment for differences in effect sizes for the association in Control versus rMDD (371/563 are FDR q < 0.05, Figure 2, Supplementary Figures 3-5).

**Figure 2:**
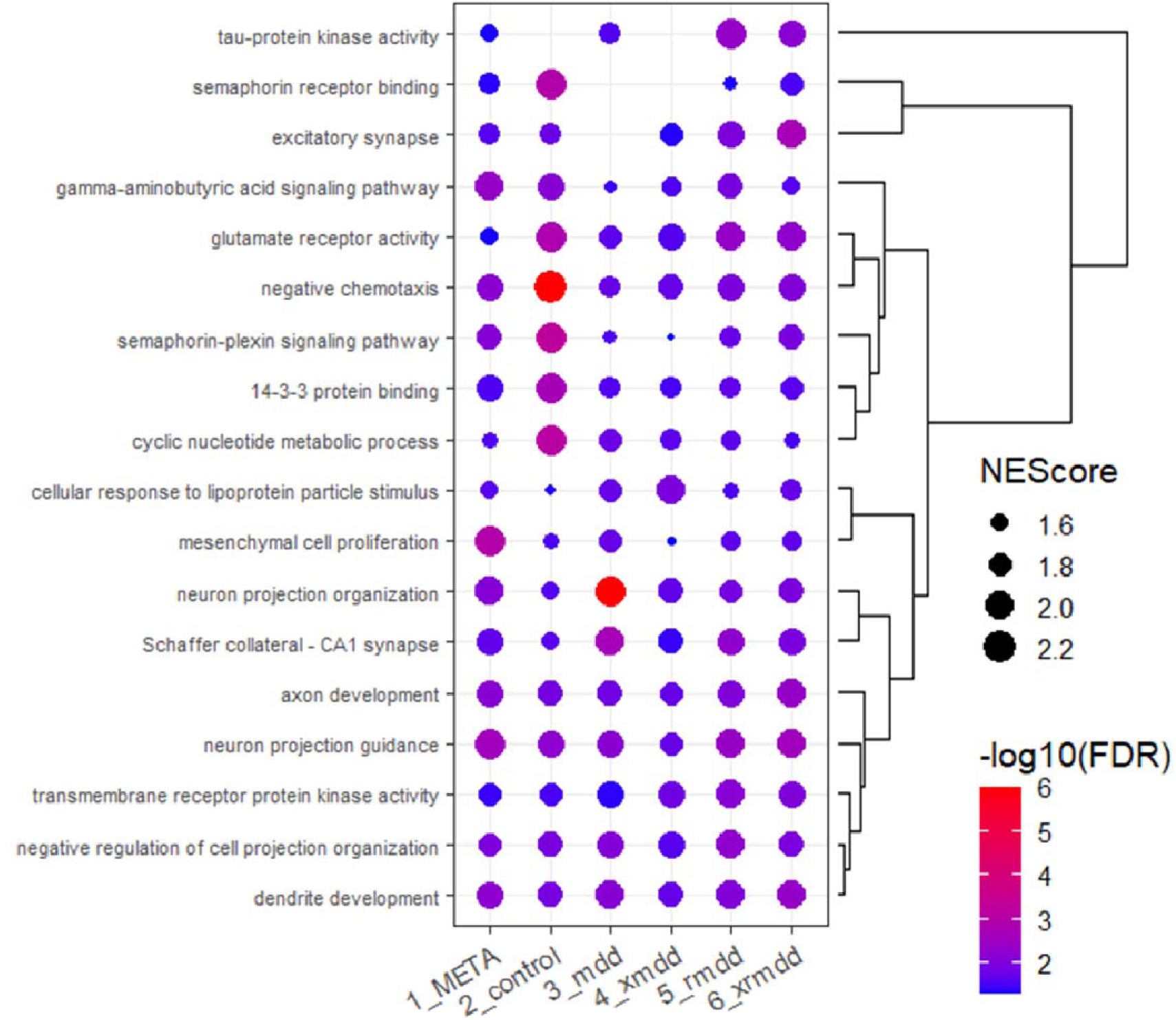
Gene set enrichment analyses of MWAS/MWEIS showing the 18 gene ontologies enrich for evidence of differential DNA methylation at an FDR q-value < 0.05 and a normalised enrichment values of > 2 in at least one group. The bubble plot point size represents the normalised enrichment score (NEScore) and the point colour represents the minus log10 of the FDR q-value. The dendrogram shows the clustering of the gene ontologies based on the NEScores using the control, MDD and rMDD.

Normalized enrichment values > 2 were found for 18 terms (Supplementary Table 15), of which 15 were FDR significant in all six MWAS/MWEIS (hypergeometric *P* = 1.7 x 10^-8^). These terms included biological processes and molecular functions associated with neuron projection development, neuron projection organization and cell surface/transmembrane receptor signalling pathways (Figure 2). The only GO cellular component term included in this set was that of Schaffer collateral - CA1 synapse. The three pathways not significant in all six MWAS/MWEIS were excitatory synapse (not FDR significant in MDD) tau-protein kinase activity (not FDR significant in Control and MDD) and semaphorin receptor binding (not FDR significant in MDD or xMDD).

There were three terms with normalized enrichment values > 2 in the META analysis. These were gamma-aminobutyric acid signaling pathway, neuron projection guidance and mesenchymal cell proliferation (FDR q-value = 0.0040, 0.0010, 0.0025, respectively). A further six terms were FDR significant in META, Control, MDD and rMDD, but not in either of the MWEIS. These were: mismatch repair, Notch signaling pathway, beta-catenin-TCF complex assembly, beta-catenin binding, hepaticobiliary system development and telomere organization. Only two terms, histone deacetylase binding and cyclase activity, were significantly enriched only in Control, MDD and rMDD, but not in the META or either MWEIS.

Twenty-two GO terms were FDR significant in the MDD-related MWAS and both MWEIS (but not in Control or META) consistent with an MDD-specific pattern. These included: regulation of neurological system process, amyloid precursor protein metabolic process, response to ischemia and one cellular component: cell division site. Twenty-six GO terms were FDR significant in four MWAS/MWEIS including META, rMDD and xrMDD, consistent with greater enrichment of putative differentially methylated genes in rMDD. These included: apical part of cell, midbrain development, neural precursor cell proliferation, semaphorin receptor binding and tau-protein kinase activity.

Of note are those terms with FDR q-value < 0.05 in all six MWAS/MWEIS analyses that contain *MATN2* or *ZEB2*. These included for *MATN2*: axon development, dendrite development, gliogenesis, neuron migration and neuron projection guidance; *ZEB2*: axon development, cell-cell signaling by WNT, central nervous system neuron differentiation, forebrain development, regulation of cell morphogenesis, regulation of neuron projection development, and regulation of protein serine/threonine kinase activity. *MATN2* containing GO terms significant in more than one group included: extracellular matrix, extracellular matrix structural constituent, regeneration, and response to axon injury. *ZEB2*-related GO terms further included, neural precursor cell proliferation, which was FDR significant in four MWAS/MWEIS: META, rMDD, xMDD, and xrMDD; and two further terms: phosphatase regulator activity and somite development that were significant in more than one analysis.

### Enrichment of MWAS/MWEIS pathways for common variants associated with adult trauma or MDD

Pathway-specific polygenic risk score analyses were performed to assess the enrichment of association signals for traumatic injury, MDD and related traits. Eighteen pathways that had normalized enrichment values > 2 and FDR q-value < 0.05 in any one MWAS/MWEIS analysis were selected (including 11 biological processes, two cellular components and five molecular functions, see Supplementary Table 15).

Polygenic scores were weighted by a GWAS for adult trauma in analysed in UK Biobank (PRS_AT,_ Supplementary Table 16; Supplementary material, Supplementary Table 17; Supplementary Figures 6-8). For PRS_AT_, neither the whole genome base set (*P* = 0.68) nor any pathway significantly predicted traumatic injury (competitive *P* > 0.05). PRS_AT_ was significantly associated with MDD (single and recurrent episode, competitive *P* = 0.0087) and recurrent MDD (competitive *P* = 0.0014) in SetA but not SetB (competitive *P* = 0.073 and *P* = 0.23, respectively). Across both SetA and SetB, enrichment for associated variants was detected for only one PRS_AT_ pathway, that of dendrite development, (competitive *P*-values SetA-rMDD = 0.040, SetB-MDD = 0.0040, SetB-rMDD = 0.024). This suggests that the dendritic development pathway is enriched for common variants associated with adult trauma that may predict lifetime MDD. Notably, this pathway is one of the three selected pathways that include *MATN2*. Four further pathway-specific PRS_AT_ were associated with MDD or rMDD in SetB: cellular response lipoprotein (MDD competitive *P* = 0.04), cyclic nucleotide metabolic process (MDD competitive *P* = 0.029 and rMDD 0.029), transmembrane receptor protein kinase activity (MDD competitive *P* = 0.012), and excitatory synapse, (rMDD competitive *P* = 0.023).

Polygenic scores were next generated for the whole genome weighted by the PGC GWAS for MDD 2018^52^ (PRS_MDD_, Supplementary Table 16). The whole genome base set significantly predicted traumatic injury (*P* = 0.0047, SetA only) and, as expected, both MDD phenotypes (P < 8.1 x 10^-26^). Enrichment for associated variants was detected for only one PRS_MDD_ pathway across both traumatic injury and both MDD phenotypes in SetA and SetB, that of the gamma-aminobutyric acid (GABA) signaling pathway, (competitive *P:* SetA-TI = 0.045, SetA-MDD = 0.010, SetA-rMDD = 0.0027, SetB-MDD = 0.016). This suggests that the GABA-signaling pathway is enriched for common variants associated with MDD. Enrichment in SetB was also seen for semaphorin receptor binding (MDD competitive *P* = 0.021); and neuron projection organisation (rMDD competitive *P* = 0.0003).

Axon development, a pathway that includes *MATN2* and *ZEB2*, was the only pathway that showed significant enrichment for the same depression phenotype, rMDD, in both the adult trauma and MDD-weighted PRS (SetA competitive *P* = 0.034, SetB competitive *P* = 0.022). Although, the best p-value threshold for PRS_AT_ was much lower, i.e. more significant, and included fewer variants, than that for PRS_MDD_.

### Gene set expression enrichment in adult human brain regions

Mapped genes from the MWAS/MWEIS were assessed for enrichment for gene expression in adult human brain regions using data from the Allen Brain Atlas. Brain regions significant at a family-wise error rate (FWER) of q < 0.05 for at least one MWAS/MWEIS are shown with the comparative results for that region in the other MWAS/MWEIS (Table 3, Supplementary Table 18). Three regions were FWER significant in the genes from MWAS of traumatic injury in individuals with rMDD: limbic lobe (MWAS/MWEIS p-values threshold (pT) = 0.005, N. genes =3 307, overlap = 1 658) and both left and right supraoptic nucleus (pT = 1 x 10^-5^, N. genes = 19, overlap = 10 genes including *MATN2*). The limbic lobe was also FWER significant in the MWEIS of MDD (pT = 0.005, N genes = 2 880, overlap=1 447).

**Table 3.** Results of regional gene-expression enrichment analyses.

| MWAS/<br>MWEIS | pT | Allen<br>ID: | Structure | FWERs for<br>expression quantiles:<br>0.5, 0.7, 0.9 | Min<br>FWER |
| --- | --- | --- | --- | --- | --- |
| **xMDD** | **2.3** | **4219** | **LL_Limbic_Lobe** | **0.027;0.543;0.602** | **0.027** |
| **rMDD** | **2.3** | **4219** | **LL_Limbic_Lobe** | **0.028;0.964;1** | **0.028** |
| MDD | 2.3 | 4219 | LL_Limbic_Lobe | 0.125;0.91;0.935 | 0.125 |
| Control | 2.3 | 4219 | LL_Limbic_Lobe | 0.605;0.992;0.453 | 0.453 |
| xrMDD | 2.3 | 4219 | LL_Limbic_Lobe | 0.595;1;0.983 | 0.595 |
| META | 2.3 | 4219 | LL_Limbic_Lobe | 0.855;0.944;0.727 | 0.727 |
| **rMDD** | **5** | **4624** | **SO_Supraoptic_Nucleus_Right** | **0.033;0.647;1** | **0.033** |
| **rMDD** | **5** | **4593** | **SO_Supraoptic_Nucleus_Left** | **0.034;0.3;1** | **0.034** |
| xrMDD | 5 | 4624 | SO_Supraoptic_Nucleus_Right | 0.581;1;0.995 | 0.581 |
| xrMDD | 5 | 4593 | SO_Supraoptic_Nucleus_Left | 0.616;0.997;0.985 | 0.616 |
| MDD | 5 | 4593 | SO_Supraoptic_Nucleus_Left | 0.66;0.992;0.96 | 0.66 |
| Control | 5 | 4593 | SO_Supraoptic_Nucleus_Left | 1;0.99;0.815 | 0.815 |
| Control | 5 | 4624 | SO_Supraoptic_Nucleus_Right | 0.972;0.99;0.877 | 0.877 |
| MDD | 5 | 4624 | SO_Supraoptic_Nucleus_Right | 0.887;0.992;0.975 | 0.887 |
| xMDD | 5 | 4593 | SO_Supraoptic_Nucleus_Left | 0.889;1;1 | 0.889 |
| xMDD | 5 | 4624 | SO_Supraoptic_Nucleus_Right | 0.984;1;1 | 0.984 |
| META | 5 | 4593 | SO_Supraoptic_Nucleus_Left | 1 | 1 |
| META | 5 | 4624 | SO_Supraoptic_Nucleus_Right | 1 | 1 |

pT = $-\log_{10}(\text{p-values})$ threshold for gene selection.

## Discussion

Trauma is associated with the onset of mental health issues such as PTSD and major depressive disorder (MDD). In this study, we investigated the association of self-reported traumatic physical injury, bone fracture occurring in traumatic circumstances, with depression and related traits. We performed a methylome-wide analysis of traumatic physical injury and DNA methylation in individuals with no mental health diagnoses and in those with major depressive disorder. Further, we compared the strength of those associations between the two groups and examined the potential functional effects through analyses of biological pathways, polygenic scores and brain regional expression.

Traumatic physical injury was associated with a lifetime diagnosis of MDD, particularly of a recurrent course, and with symptoms associated with psychiatric illness. There was no association with cognitive traits, apart from with digit symbol coding score, a measure of processing speed which is associated with schizophrenia spectrum disorders.^50, 51^ The strongest association was seen with the SPQ cognitive disorganization factor, which is characterised by deficits in ability to organize and express thoughts and behaviour. This subscale is strongly positively associated with an individual’s personal and familial psychiatric history.^35^ Childhood trauma score has previously been associated with both SPQ total score and subscales.^53^ A thought disorder factor has previously been associated with poor mental health, particularly physical abuse and intrusive memories following trauma.^54-58^

Methylome-wide analyses identified opposing patterns of DNA methylation, on average, when comparing the MDD and control groups, we identified when examining the approximately 40,000 sites nominally associated with traumatic injury. This pattern was most prominent at the first exon and exon boundaries, despite a significant underrepresentation of differentially methylated sites at the 1st exon in the MDD group. Although, the association between levels of DNA methylation and gene expression are difficult to predict on such a wide scale, this may reflect a reduction in gene expression and/or increased alternative splicing at key genes.^59, 60^

Examination of the DNA methylation profiles associated with traumatic injury identified two CpG sites with genome-wide significant differential DNA methylation: cg14764459 (Control *P* = 5.14 x 10^-8^, META *P* = 6.76 x 10^-8^) and cg02101279 (rMDD *P* = 8.60 x 10^-8^). Although cg14764459 maps to LOC100287944, it lies in an ENCODE distal enhancer-like signature (cCRE ENCODE Accession: EH38E1641208) linked to *RIC8B* and containing a putative glucocorticoid response element.^61^ RIC8B is a non-canonical GEF for heterotrimeric G protein alpha subunits. It is expressed in neural crest cells, astrocytes, Purkinje cells, the cerebellum, hippocampus and adrenal gland, is associated with neurodevelopment, cortical surface area, cortical thickness, and may regulate synapse formation.^62, 63^ The second genome-wide significant CpG site, cg02101279 (rMDD *P* = 8.60 x 10^-8^), is also associated with a distal enhancer-like sequence with NR3C1-binding potential (EH38E2155469). However, the CpG site is six base-pairs from the enhancer and there is no gene linked to this sequence in the ENCODE SCREEN database.^61^

Downstream analyses of the MWAS/MWEIS showed significant overlap in the gene ontology pathways enriched for signals of differential methylation associated with physical trauma in individuals with and without MDD. Although this initial study suggests that epigenetic response to traumatic injury may be associated with the same functional processes in individuals regardless of their mental health diagnoses; the magnitude, and indeed the direction, of the association may differ at the levels of individual CpG sites, genes and functional pathways. Gene set enrichment analyses identified enriched DNA methylation signals in key neuronal development and neuronal signalling pathways including axon development, dendrite development and GABA-ergic signalling suggesting possible functional effects on synaptic plasticity.^64, 65^ These pathways included *MATN2* and/or *ZEB2,* genes linked to CpG sites with suggestive levels of differential methylation in MDD groups (see: Supplementary Note – *MATN2* & *ZEB2*), and were enriched for common variants associated with MDD and adult trauma. GABA-ergic signalling has previously been associated with depression^66^ and trauma, particularly involving the limbic system.^67-70^ and may be important in the response to cognitive therapy targeting the suppression of traumatic memories.^71^ A recent study of depression in twins also identified glutamate and GABA-ergic, as well as ECM-receptor interaction, as enriched in a weighted-gene co-expression network cluster associated with depressive symptoms and disease state.^72^ This study also highlighted the regulation of NOTCH-signalling, a pathway strongly associated with *MATN2*.

Finally, regional brain enrichment analyses suggested that the genes mapping to CpG sites with differential DNA methylation *P* < 1 x 10^-5^ in individuals with rMDD were enriched for genes expressed in the limbic lobe (Allen Brain Atlas ID: 4219, containing the cingulate gyrus, hippocampal formation, parahippocampus, and piriform cortex) and supraoptic nuclei (Allen Brain Atlas ID: 4593, 4624; part of the anterior hypothalamic region). This is consistent with the enrichment of gene ontology term Schaffer collateral - CA1 synapse in all the MWAS/MWEIS analyses. Lesions of hippocampal CA1 neurons significantly effects retrieval of episodic memories^73, 74^ and these neurons are important for fear extinction.^75^ The limbic and hippocampal regions have previously been linked to anxiety, fear memory formation and PTSD.^67^ The supraoptic nuclei contain vasopressin and oxytocin-producing cells and has been implicated in responses to acute restraint.^76^ Vasopressinergic neurons in the SON sending projections to limbic forebrain structures such as the dorsal hippocampus, ventral hippocampus, BNST, and amygdala,^77^ and can control hippocampal excitability and response to acute restraint stress.^76^

This study has limitations, the restriction of these analyses to white British individuals only, reduces the representativeness of the results. The use of Generation Scotland participants may also have resulted in the individuals having less severe depression, higher income, and higher cognitive ability on average than in the general population, as is common for population-based samples. A major limitation of this study is the sample size, particularly of the MDD groups, replication is required in larger samples. The trauma weighted PRS analyses was limited by the availability of a suitable adult trauma genome-wide association study. We used a previously defined adult trauma phenotype, but this phenotype may differ from the traumatic physical injury phenotype. This study examined DNA methylation in whole blood and, although whole blood has shown some correlation with DNA methylation changes in the brain particularly those sites under genetic control, this may reduce the power to detect regional or cell types important to response to traumatic injury. Analysis of whole blood increases the available sample size and may provide a biomarker to guide treatment and/or for treatment response.^78^ Further, we were unable to examine the temporal association of the phenotypes. The traumatic physical injury may have occurred before or after the onset of major depression and current diagnostic symptoms were not included as covariates in the analyses. The differences in DNA methylation levels may therefore reflect genetic susceptibility loci, whist others may reflect the disease state or medication. The identification of differential methylation in those individuals without a diagnosis of mental illness may, however, suggest a direct association with traumatic injury. However, the collection of samples pre- and post-event would be required to distinguish between potential causal mechanisms.

In conclusion, these results suggest that traumatic physical injury is associated with widespread differential DNA methylation. Particularly associated with genes involved in synaptic plasticity and glutamate and GABA-ergic signalling pathways, and in regions of the brain associated with encoding fear memory. However, these results require replication in larger cohorts and the use of longitudinal study designs. Such studies may help to identify markers for the long-term effects of traumatic injury, particularly on mental health, and identify therapies and treatments.

## Acknowledgements

Ethics approval for Generation Scotland was given by the NHS Tayside committee on research ethics (reference 05/S1401/89). Generation Scotland received core funding from the Chief Scientist Office of the Scottish Government Health Directorate CZD/16/6 and the Scottish Funding Council HR03006. Genotyping of the GS:SFHS samples was carried out by the Genetics Core Laboratory at the Wellcome Trust Clinical Research Facility, Edinburgh, Scotland and was funded by the UK’s Medical Research Council and the Wellcome Trust (Wellcome Trust Strategic Award “STratifying Resilience and Depression Longitudinally” (STRADL) (Reference 104036/Z/14/Z). This work has made use of the resources provided by the Edinburgh Compute and Data Facility (ECDF). (http://www.ecdf.ed.ac.uk/).

The research was conducted using the UK Biobank resource - application number 4844. UK Biobank received ethical approval from the North West Multi-centre Research Ethics Committee in the UK (reference 11/NW/0382), and the current study received approval from the UKB Access Committee (application #4844). All participants gave written informed consent.

We are grateful to all the families who took part, the general practitioners and the Scottish School of Primary Care for their help in recruiting them, and the whole Generation Scotland team, which includes interviewers, computer and laboratory technicians, clerical workers, research scientists, volunteers, managers, receptionists, healthcare assistants and nurses.

