## Supplementary Material for "Methylome-wide association studies of traumatic injury identifies differential DNA methylation of synaptic plasticity and GABAergic-signalling"

Jodie Brown *et al.*

**Supplementary Notes**

**Genome-wide Association of Adult Trauma	Page #**

Adult Trauma Phenotype UK Biobank	2

Genome-wide association study	3

Results	4

***MATN2* & *ZEB2***	5

**Supplementary References**	7

**Supplementary Tables**

See: Supplementary tables MWAS traumatic injury.xls

**Supplementary Figures**

Supplementary Figure 1a-f. Manhattan plots of the MWAS and MWEIS analyses with methylome-wide significant CpG sites annotated (p-value < 9.42 x 10^-8^)	10

Supplementary Figure 2. The overlap between genes mapped to CpG sites with p-values < 1x10^-5^	11

Supplementary Figure 3: Overlap of the gene ontologies in enriched in the MWAS analyses at FDR q-values < 0.05	12

Supplementary Figure 4: Overlap of the gene ontologies in enriched in the MWAS/MWEIS analyses of Control and rMDD at FDR q < 0.05.	13

Supplementary Figure 5: Overlap of the gene ontologies in enriched in the MWAS/ analyses of Control and rMDD at FDR q < 0.05.	13

Supplementary Figure 6: SNP and gene-based Manhattan plots of the adult trauma GWAS in UK Biobank	14

Supplementary Figure 7: MAGMA results for adult trauma GWAS in UK Biobank	15

Supplementary Figure 8: GTEx enrichment results for adult trauma GWAS in UK Biobank	16

**Supplementary Note: Genome-wide Association of Adult Trauma**

Adult Trauma Phenotype UK Biobank

The mental health questionnaire (MHQ) was developed with the intention of phenotyping mental health within UK Biobank ^1^. The MHQ took the form of an online questionnaire and has accrued upwards of 150 000 responses. It utilises a variety of already validated measures, such as the Composite International Diagnostic Interview Short Form (CIDI-SF, Kessler et al. 1998), used to identify MDD. MDD and adult trauma data from white British individuals was derived from the MHQ to minimise the impact of population stratification on GWAS results.

The adult trauma phenotype was derived as part of this project using UK Biobank data in R v4.2.2. Case definitions for adult trauma within UK Biobank applied following: ^2^ Adult trauma cases and controls were categorised using the MHQ responses in fields 20522-20525. For each question, those who selected a response indicating that they have experienced that specific trauma were considered cases and were coded as “1”. Those who did not report that they had experienced the described trauma were controls and coded as “0”. Those who responded with “Prefer not to answer” were excluded from analysis. This resulted in each individual having a trauma score between 0 and 5, based on the number of adult traumas they had experienced.

UK Biobank Mental health questionnaire (MHQ) Adult trauma fields.

| Field ID | Question  “Since I was sixteen… | Response options | Adult trauma criteria |
| --- | --- | --- | --- |
| 20522 | …I have been in a confiding relationship” | “Never true”  “Rarely true”  “Sometimes true”  “Often”  “Very often true”,  “Prefer not to answer”. | Responses of  “Never true”, “Rarely true”, and “Sometimes true”  were all considered cases |
| 20523 | …a partner or ex-partner deliberately hit me or used violence in any other way” |  | Responses of  “Rarely true”, “Sometimes true”, “Often true”, and “Very often true”  were all considered cases |
| 20521 | …a partner or ex-partner repeatedly belittled me to the extent that I felt worthless” |  | Responses of  “Rarely true”, “Sometimes true”, “Often true”, and “Very often true”  were all considered cases |
| 20524 | …a partner or ex-partner sexually interfered with me, or forced me to have sex against my wishes” |  | Responses of  “Rarely true”, “Sometimes true”, “Often true”, and “Very often true”  were all considered cases |
| 20525 | …there was money to pay the rent or mortgage when I needed it” |  | Responses of  “Never true”, “Rarely true”, “Sometimes true”, and “Often true”  were all considered cases |

Genome-wide association study

Unrelated UK white British individuals were selected from UK Biobank as in Howard *et al.,* briefly, participants were removed based on UKB genomic analysis exclusion, non-white British ancestry, high missingness, genetic relatedness (kinship coefficient > 0.0442), QC failure in UK BiLEVE study and gender mismatch. UKB received ethical approval from the NHS National Research Ethics Service North West (reference: 11/NW/0382). Further details on UKB cohort description, genotyping, imputation and quality control are available elsewhere.^3, 4^

Adult trauma score was analysed as a continuous variable using linear regression under an additive model using Pink V1.9b4 ^5^. Age, sex, and 20 genetic principal components were fitted as covariates to account for cryptic population stratification and known confounders. Individuals with incomplete phenotypic data were excluded from analysis. The final data set was comprised of 76 796 individuals. 26 654 of which 37 699 were adult trauma cases. The distribution of sex is roughly even, with 54% of the sample being female, and 46% male. UK Biobank participants were aged between 37- to 73-years at recruitment in 2014. As a result of this, the age range of the samples used in this study is 46.5- to 80.5-years, with a median age of 65.1-years. Female participants were more likely to report traumatic events in adulthood than male, at 56% and 45% respectively. All ages had comparable distribution of adult trauma cases and control.

MAGMA, within FUMA, was used to perform gene-based tests using the GWAS summary statistics, mapping input SNPs to protein coding genes ^6^. SNPs were mapped to genes when they were within 10kb. From this, FUMA conducts analysis of gene expression by tissue, tissue specificity for mapped genes using; ‘GTEx v8: 54 tissue types’ and ‘GTEx v8: 30 tissue types’, and gene-set enrichment analysis. The significance threshold used was a Bonferroni corrected *P*-value of < 0.05. The major histocompatibility complex located in chromosome 6p21 was excluded from analysis.

Two p-value thresholds were used: *P* < 5x10^-8^ represents genome-wide significance, and *P* < 1x10^-5^ represents suggestive significance and better captures alleles that occurs at a lower frequency.

Results

The GWAS of adult trauma tested 552 228 genetic variants and 74 224 individuals. No results were genome-wide significance. Eleven individual SNPs reached suggestive significance (see below) and no genes were significant in the gene-based test Q-Q plots show minor inflation, lambda value, 1.108. Seven lead SNPs at suggestive significance mapped to 16 genes. See: Supplementary Figure 10.

Eight mapped genes are highly expressed in brain tissues and an additional 22 mapped genes are highly expressed across numerous tissues as well as brain tissues (Supplementary Figure 11).

No tissue type reached significance after correction for multiple testing in MAGMA tissue expression analysis. However, the pituitary, stomach, and EBV-transformed lymphocytes were nominally significant (*P* < 0.05) for ‘GTEx v8 54 general tissue types’, (Supplementary Figures 10-12).

SNPs reaching suggestive significance levels:

| SNP | *P* | Chr | Mapped genes |
| --- | --- | --- | --- |
| rs10432523 | 2.613x10^-6^ | 2 | *PRPF40A; ARL6IP6; RPRM; GALNT13; KCNJ3; NR4A2; GPD2* |
| **rs4664759** | 1.789x10^-6^ | 2 | *PRPF40A; ARL6IP6; RPRM; GALNT13; KCNJ3; NR4A2; GPD2; AC011308.1; GALNT5; ERMN; CYTIP; ACVR1C* |
| rs1900132 | 8.539x10^-6^ | 2 | *RPRM; GALNT13; KCNJ3; GPD2; GALNT5; ACVR1C* |
| rs13110423 | 4.457x10^-6^ | 4 | *FBXL5; FAM200B; CD38; TAPT1; LDB2; QDPR; CLRN2* |
| **rs283019** | 1.80x10e^-6^ | 4 | *FBXL5; FAM200B; CD38; TAPT1; LDB2; QDPR; CLRN2* |
| **rs10477349** | 3.647x10^-6^ | 5 | *GPR151; DPYSL3; JAKMIP2; SPINK1; HTR4; SH3TC2* |
| rs2466432 | 2.753x10^-6^ | 8 | *RAD54B; FSBP; KIAA1429; AC023632.1; ESRP1; INTS8; CCNE2; TP53INP1; PLEKHF2; C8orf37; UQCRB; MTERFD1; PTDSS1; NDUFAF6* |
| **rs16916994** | 1.009x10^-6^ | 8 | *RAD54B; FSBP; KIAA1429; ESRP1; DPY19L4; INTS8; CCNE2; TP53INP1; C8orf37; UQCRB; MTERFD1; PTDSS1; NDUFAF6* |
| **rs10787532** | 2.141x10^-7^ | 10 | *MAT1A; DYDC1; DYDC2; FAM213A; SH2D4B; NRG3; GHITM* |
| **rs4760663** | 4.437x10^-6^ | 12 | *LINC00935; ADCY6; CACNB3; DDX23; RND1; RP11-302B13.5; CCDC65; FKBP11; AC073610.5; ARF3; WNT10B; WNT1; DDN; PRKAG1; KMT2D; DHH; TROAP; C1QL4 SPATS2* |
| **rs4799903** | 7.935x10^-7^ | 18 | *DSC3* |

Bold = lead SNPs *P* < 1x10^-5^

**Supplementary Note – *MATN2* & *ZEB2***

Sixty-six CpG sites showed differential DNA methylation at the suggestive threshold of *P* < 1 x 10^-5^. Of these, two CpG sites, mapping to genes *MATN2* and *ZEB2*, were associated with differential methylation in the analyses of individuals with MDD with evidence for greater differential methylation in individuals with MDD than controls (MWEIS p < 1 x 10-5). MATN2 encodes the protein Matrilin-2, a component of the extracellular matrix important for mediating interactions in collagen-dependent and collagen-independent filamentous networks (Piecha et al. 1999) Common variants in *MATN2* have previously been associated with the interaction between childhood trauma events and smoking behaviour^7^. *MATN2* variants are also associated with MATN2 and NELL2 (Neural EGFL-Like 2) protein levels (Consortium 2013). NELL2 is thought to be involved in nervous system development and regulation of SLIT/ROBO2 signalling in axon guidance ^8^. MATN2 is expressed in the brain in the extracellular matrix^9^. Extracellular matrices, in particular perineuronal nets, regulate synaptic plasticity and may have a role in memory formation and fear extinction ^10-13^. Expression of MATN2 may increase neuroinflammation in response to infection ^14^. Increased expression of MATN2 has previously been reported in the post-mortem amygdala samples from men with MDD, particularly in those individuals whose expression profiles were positively correlated with those of an unpredictable chronic mild stress mouse model of depression ^15^. This pattern was absent in individuals showing remittance or partial remittance at time of death. Interestingly, MATN2 expression in the amygdala was reduced on treatment of the mouse model with antidepressants targeting serotonergic or neuroendocrine stress pathways (Fluoxetine or corticotropin-releasing-factor 1 antagonist SSR125543). The amygdala is important for long-term storage of highly emotive and fear-related memories. Valproic acid has also been shown to decrease MATN2 expression ^16-20^.

ZEB2 (zinc finger E-box binding homeobox 2, previously known as ZFHX1B) encodes a negative regulator of DNA transcription. Genetic variants in ZEB2 are associated with smoking initiation, schizophrenia, educational attainment, risk taking behaviour and Mowat-Wilson Syndrome (featuring microcephaly, mental retardation, epilepsy, and characteristic facial dysmorphia). ZEB2 expression inhibits integrin signalling, reducing neuronal adhesion to the extracellular matrix during development to ensure the correct organisation of excitatory neurons and cortical interneurons ^21, 22^. Valproic acid has been shown to increase ZEB2 expression and is the most commonly prescribed, and effective, antiepileptic drug for Mowat-Wilson Syndrome (reviewed in ^23^).

The potential regulation of both *MATN2* and *ZEB2* expression by valproic acid (VPA) may indicate a therapeutic role for histone deacetylase inihibitors (HDACi) for psychiatric symptoms resulting from traumatic injury.^24^ VPA is a relatively nonselective HDAC Classes I and II inhibitor, known to enhance histone acetylation to promote synaptic plasticity, thereby facilitating fear extinction learning.^25^ VPA may increase GABAergic transmission, and normalise glutamate neurotransmission.^26, 27^ VPA also upregulates NOTCH1 signalling, at least in proliferating cells. In a large animal brain injury model VPA treatment reduced injury and improved recovery,^28^ but trials in humans indicate heterogeneity in response to HDAC inhibition.^29^ Evidence, although limited, suggests that use of VPA in conjunction with cognitive behaviour therapy may reduce symptoms of panic disorders and improve fear extinction.^30-32^ Roseberry et al, suggest that Valproate could provide a possible therapeutic route for anxiety disorders.^33^ This is consistent with VPA, prescribed for the treatment of bipolar disorder, as a possible therapeutic approach to trauma, particularly for irritability associated with trauma.^34^

**Supplementary Figure 1.** Manhattan plots of the MWAS and MWEIS analyses with methylome-wide significant CpG sites annotated (p-value < 9.42 x 10^-8^). Blue horizontal line represents p-value = 1 x 10^-5^

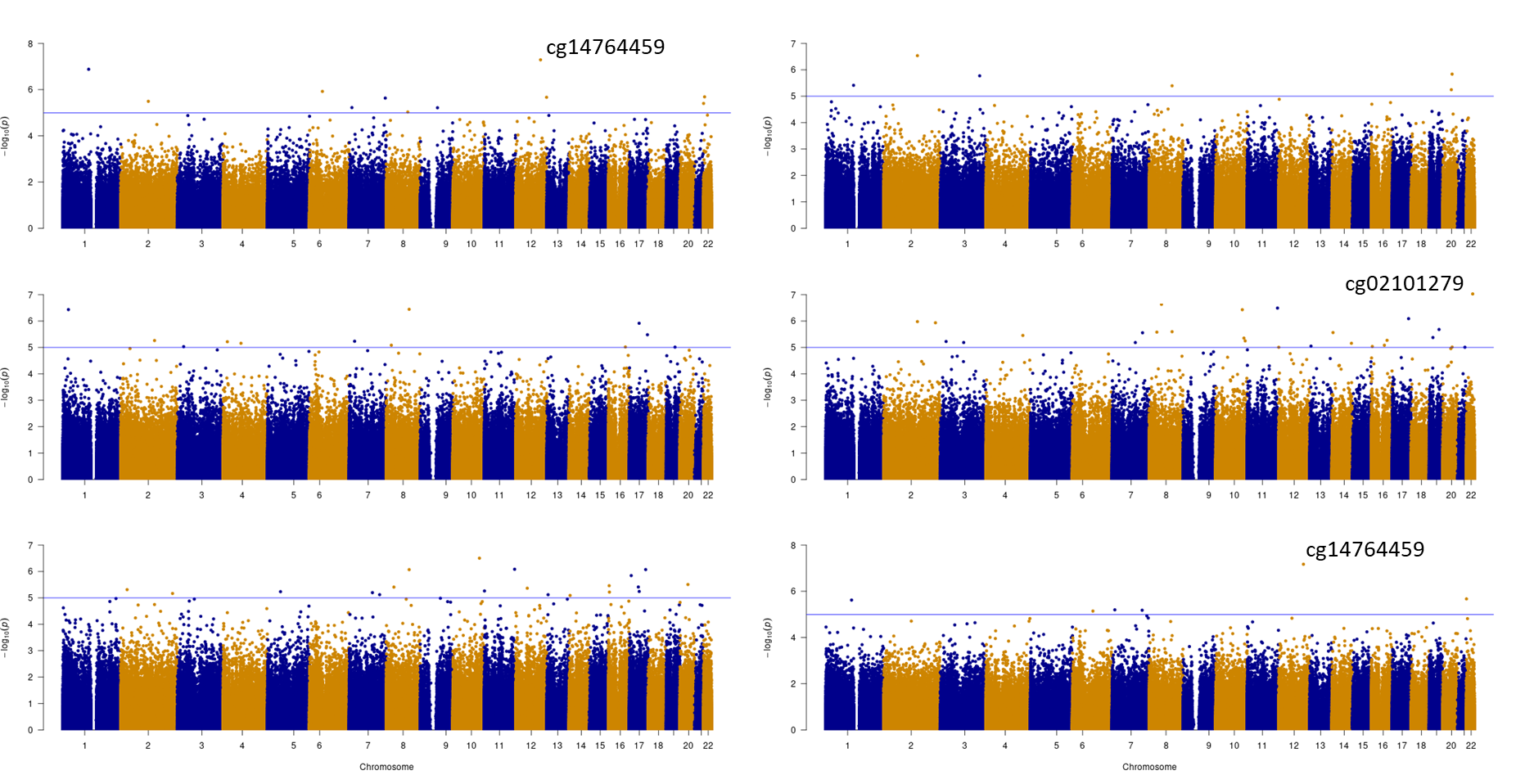
a) MWAS Control	d) MWEIS xMDD

b) MWAS MDD	e) MWEIS xrMDD

c) MWAS rMDD	f) MWEIS META

**Supplementary Figure 2.** The overlap between genes mapped to CpG sites with p-values < 1x10^-5^.

Venn diagrams of the MWAS results and tables of the MWAS genes

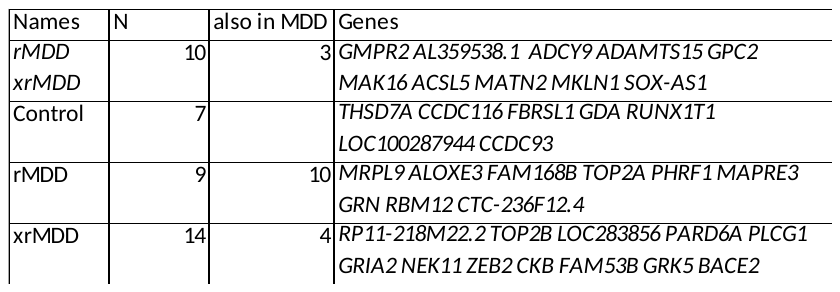

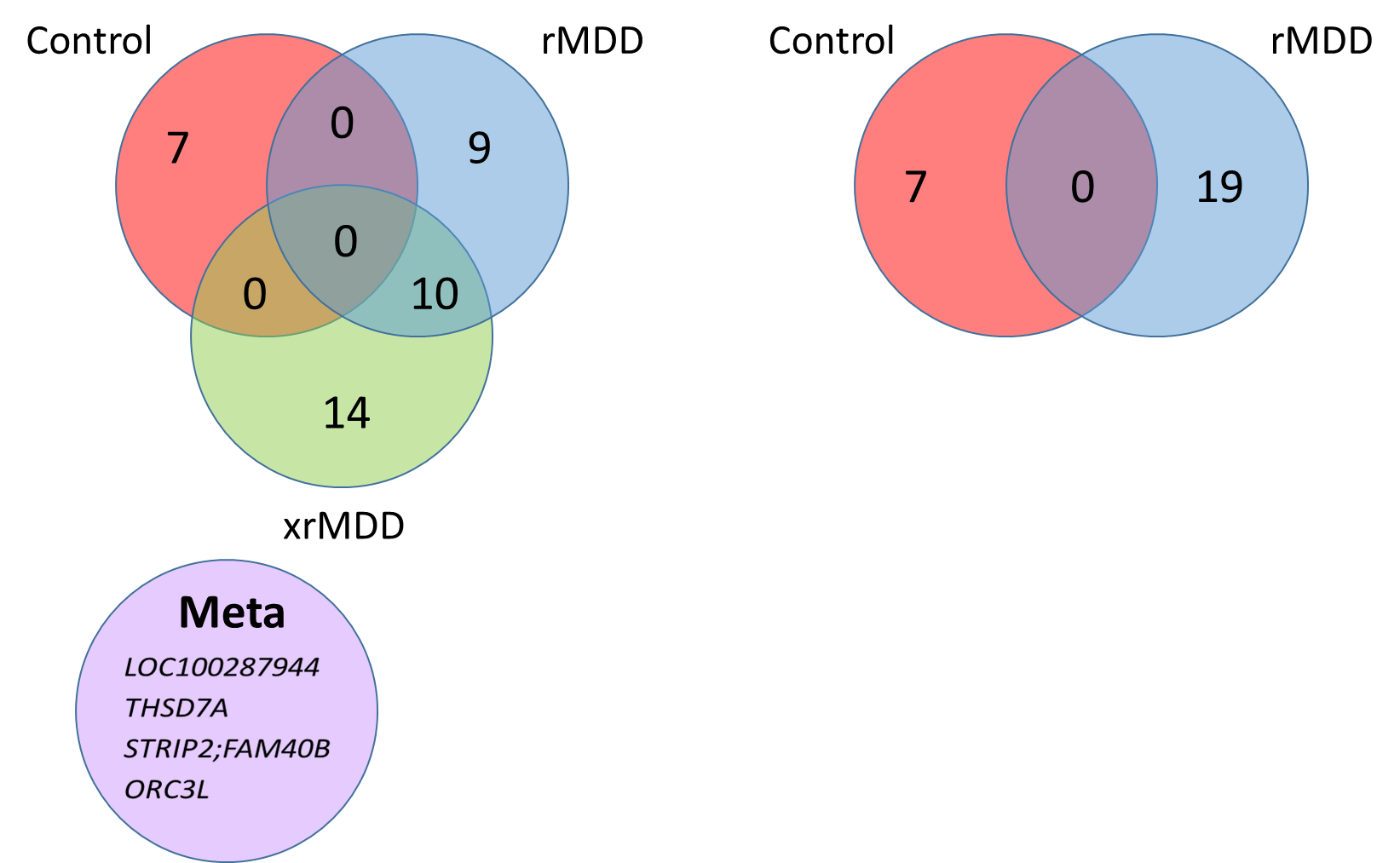

**Supplementary Figure 3.** Overlap of the gene ontologies in enriched in the MWAS analyses

at FDR q-values < 0.05 (out of 10,532 ontologies tested).

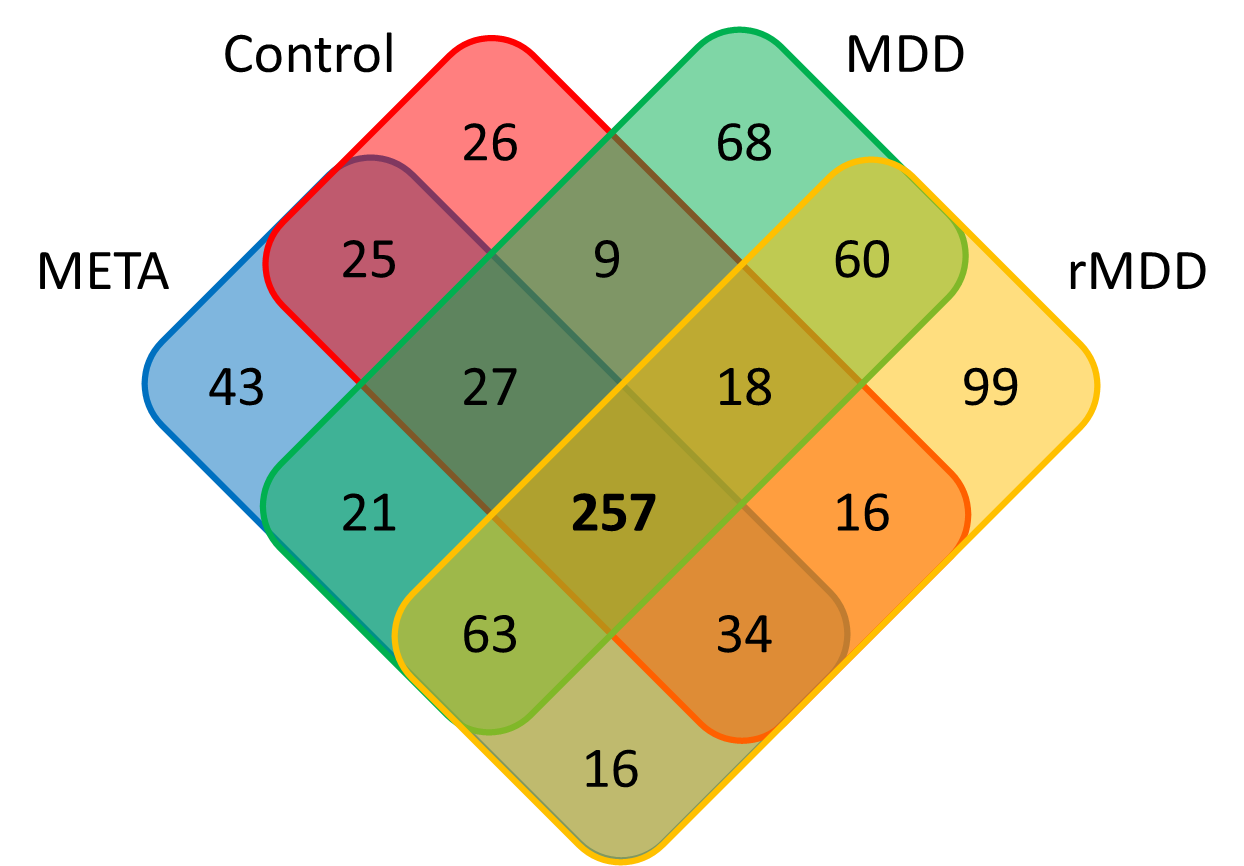

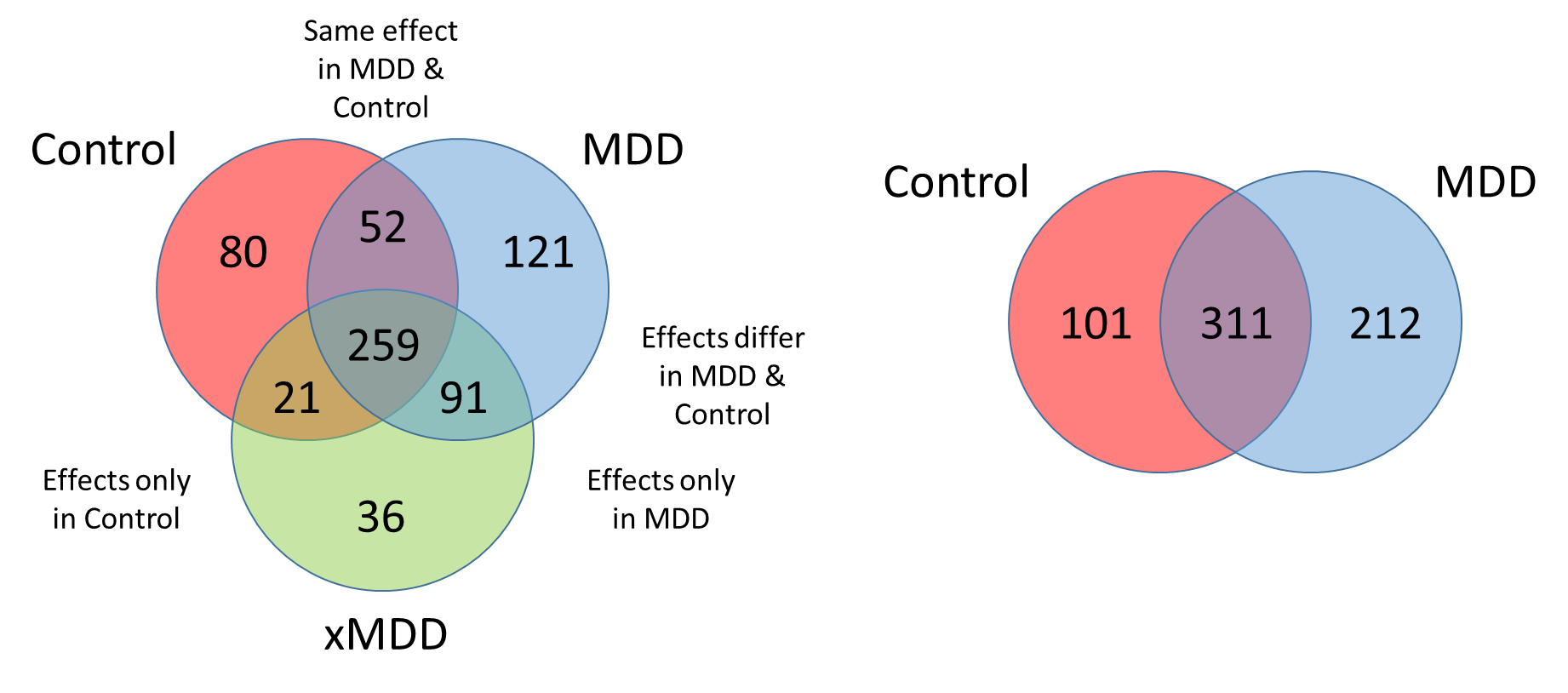
**Supplementary Figure 4.** Overlap of the gene ontologies in enriched in the MWAS/MWEIS analyses of Control and MDD at FDR q < 0.05.

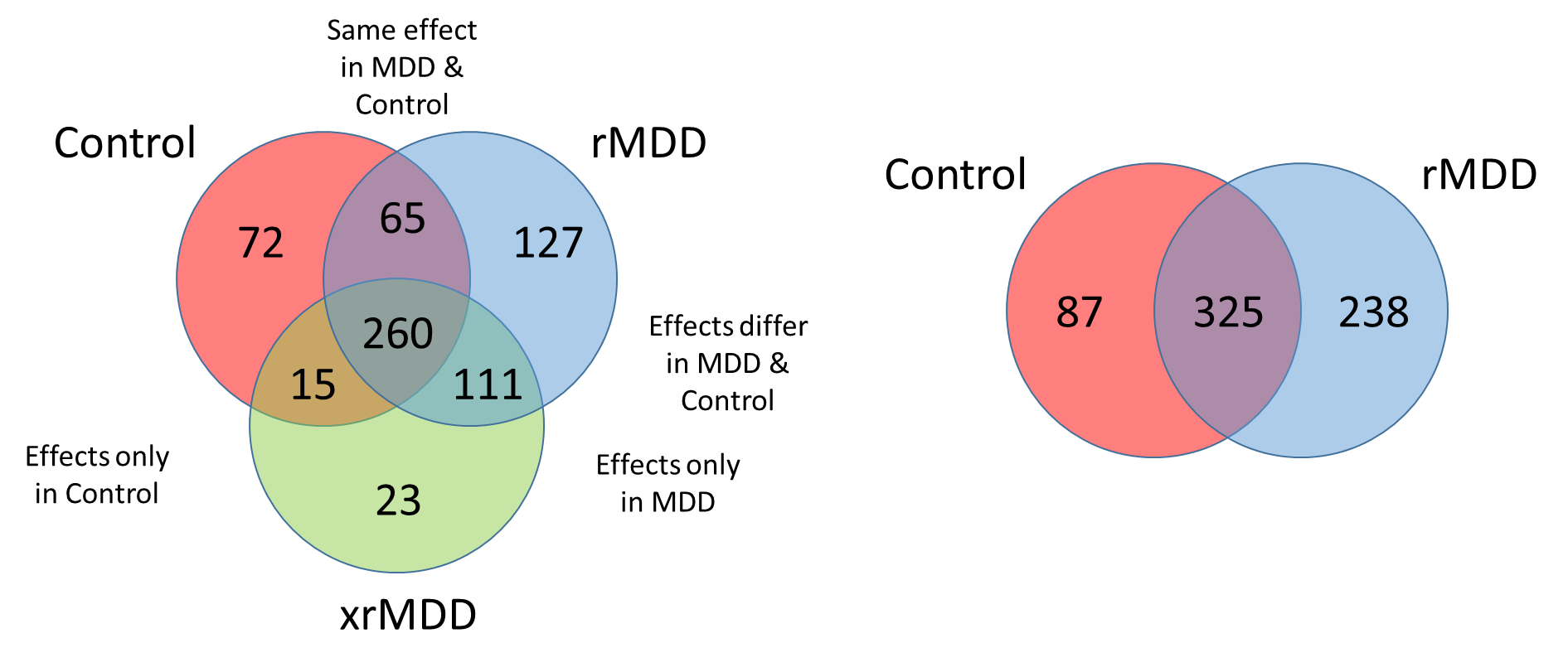
**Supplementary Figure 5.** Overlap of the gene ontologies in enriched in the MWAS/MWEIS analyses of Control and rMDD at FDR q < 0.05.

**Supplementary Figure 6.** SNP (top) and gene-based (bottom) Manhattan and QQ-plots of the adult trauma GWAS in UK Biobank

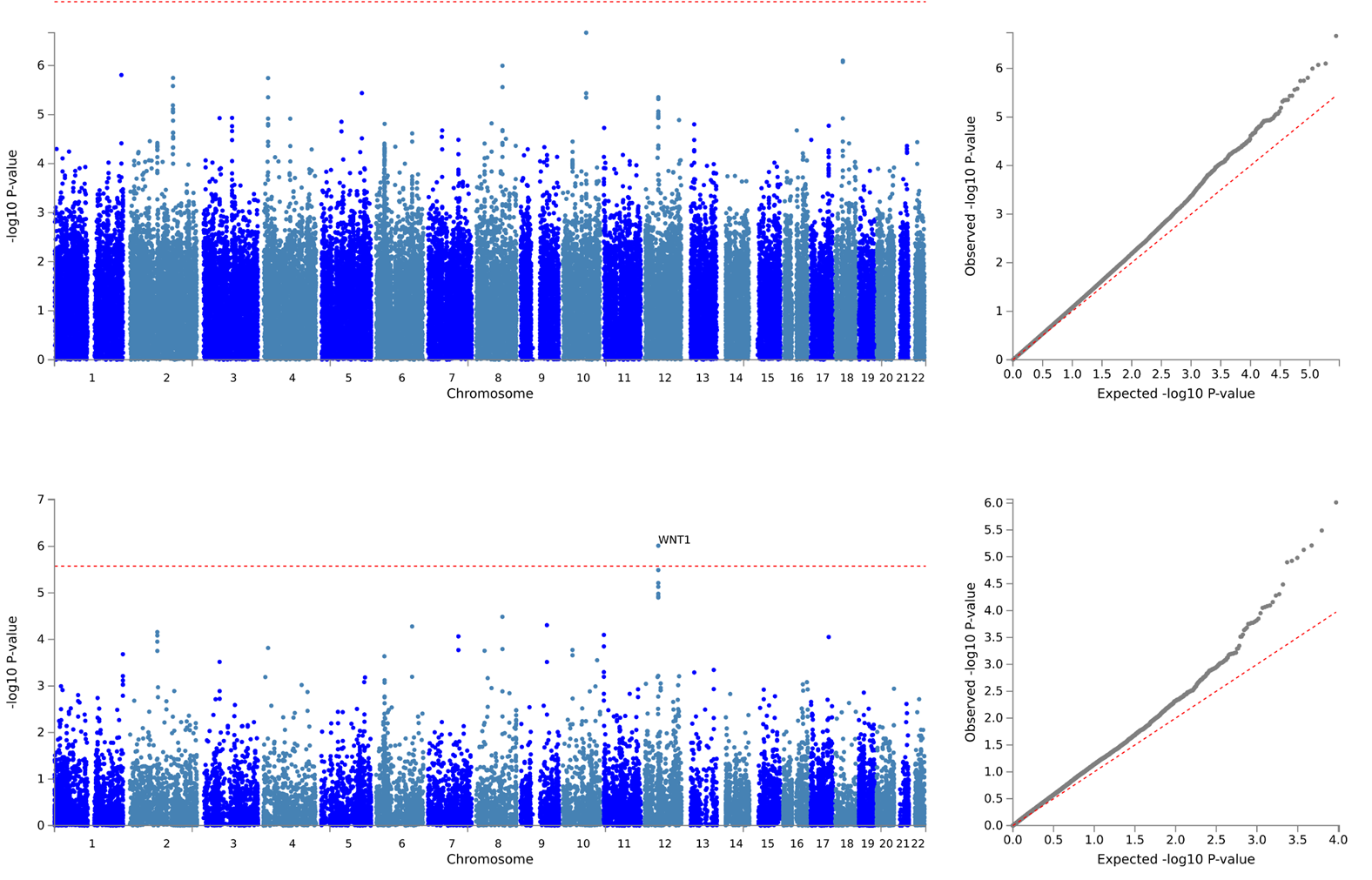

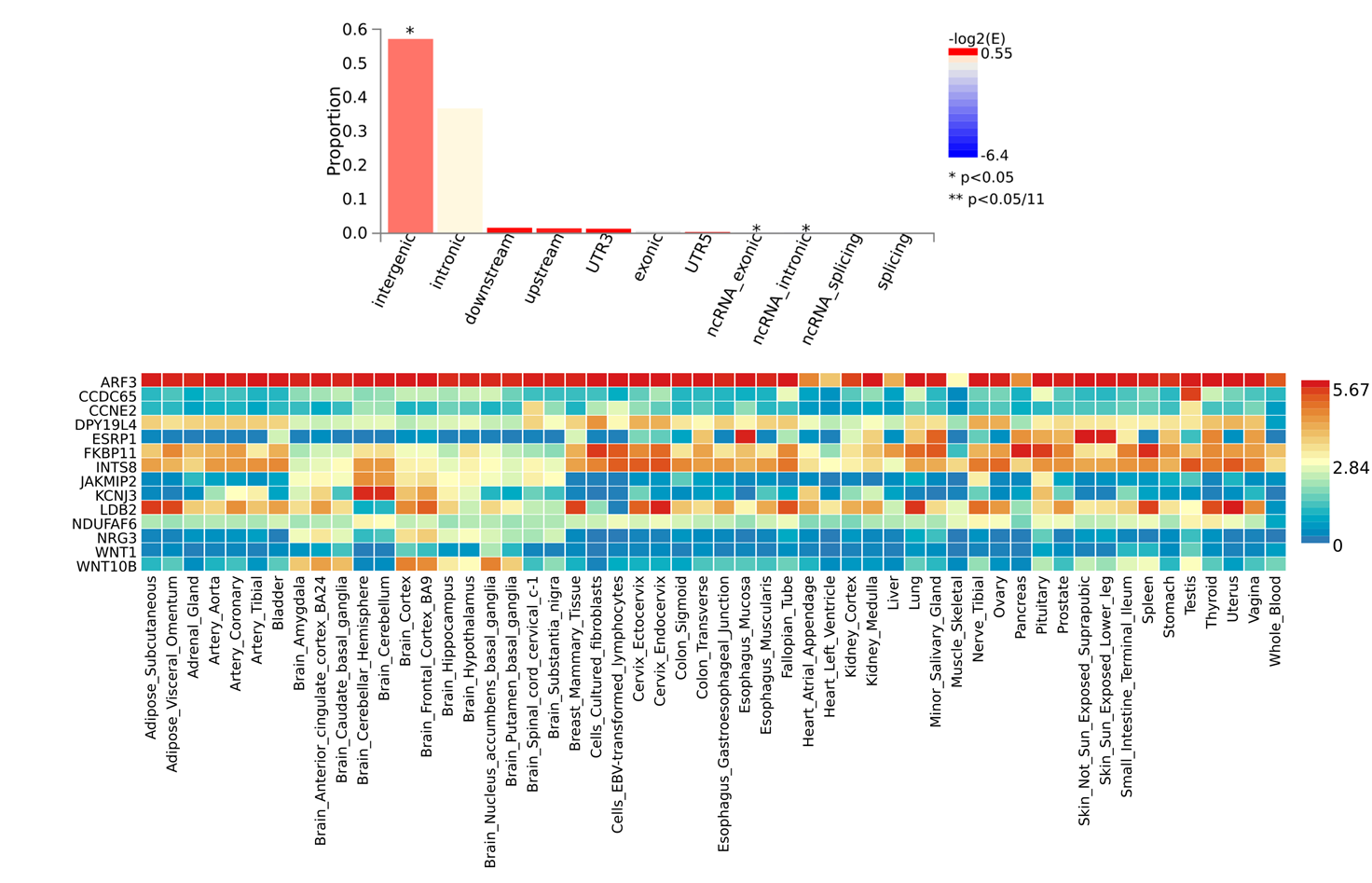
**Supplementary Figure 7.** MAGMA results for adult trauma GWAS in UK Biobank

**Supplementary Figure 8.** GTEx enrichment results for adult trauma GWAS in UK Biobank

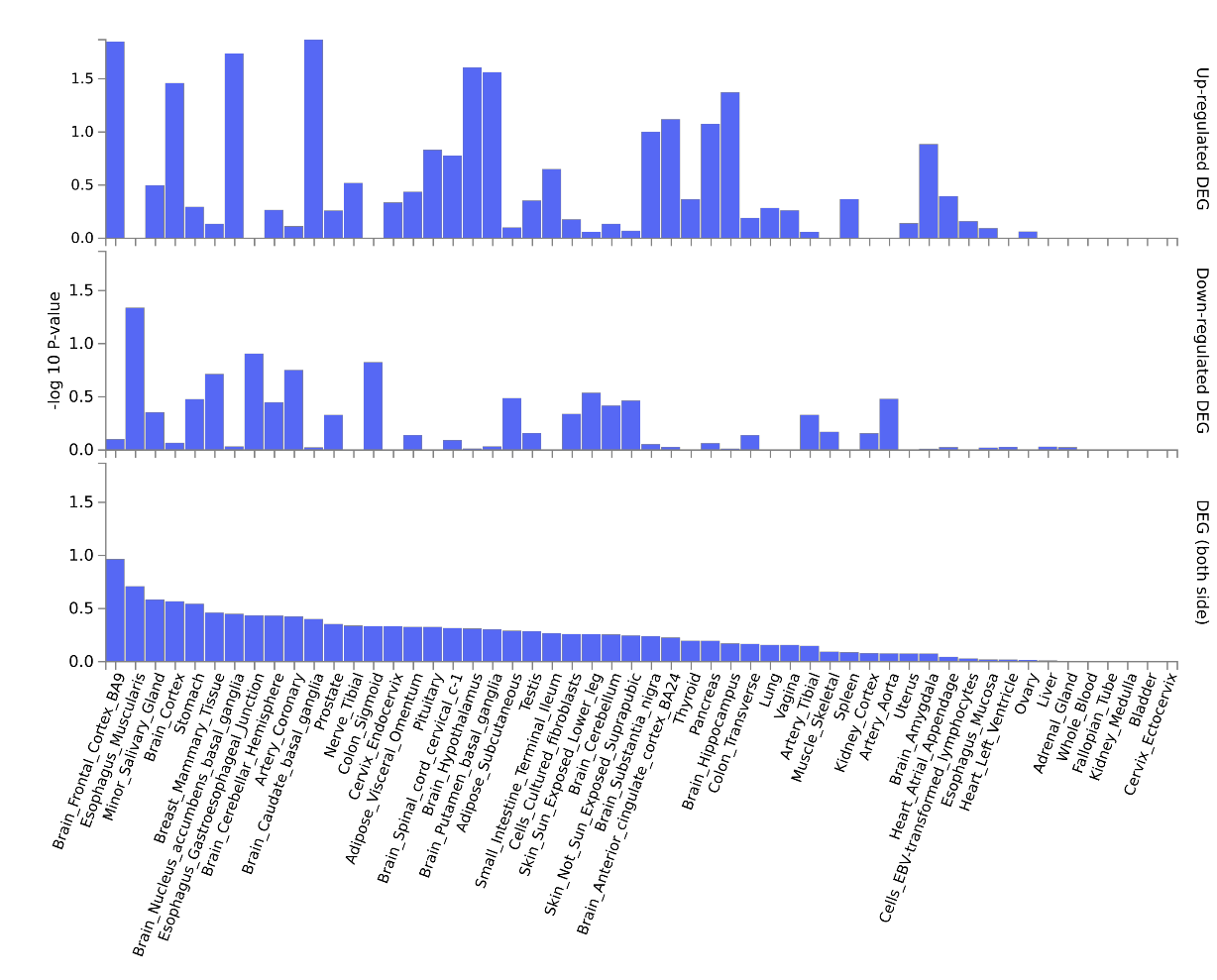

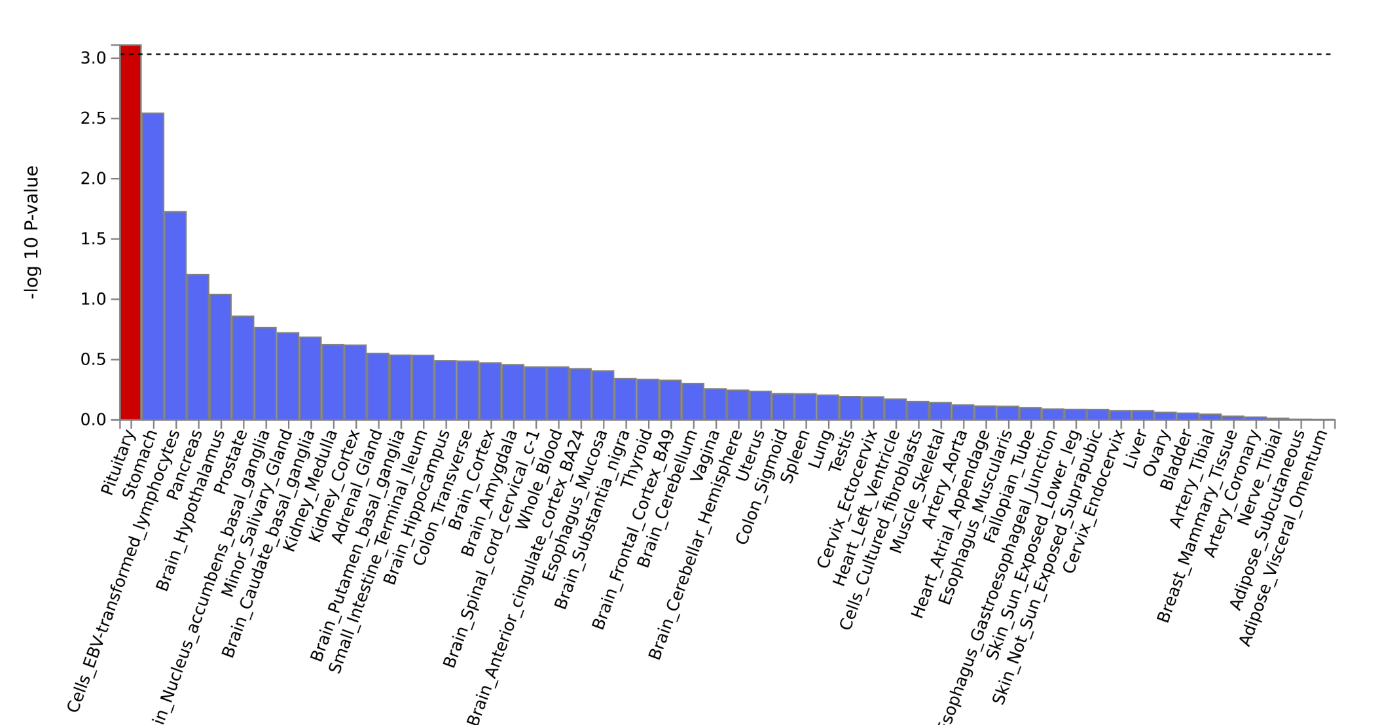
