## Supplementary Figures for "Methylome-wide association studies of traumatic injury identifies differential DNA methylation of synaptic plasticity and GABAergic-signalling"

### Slide 1
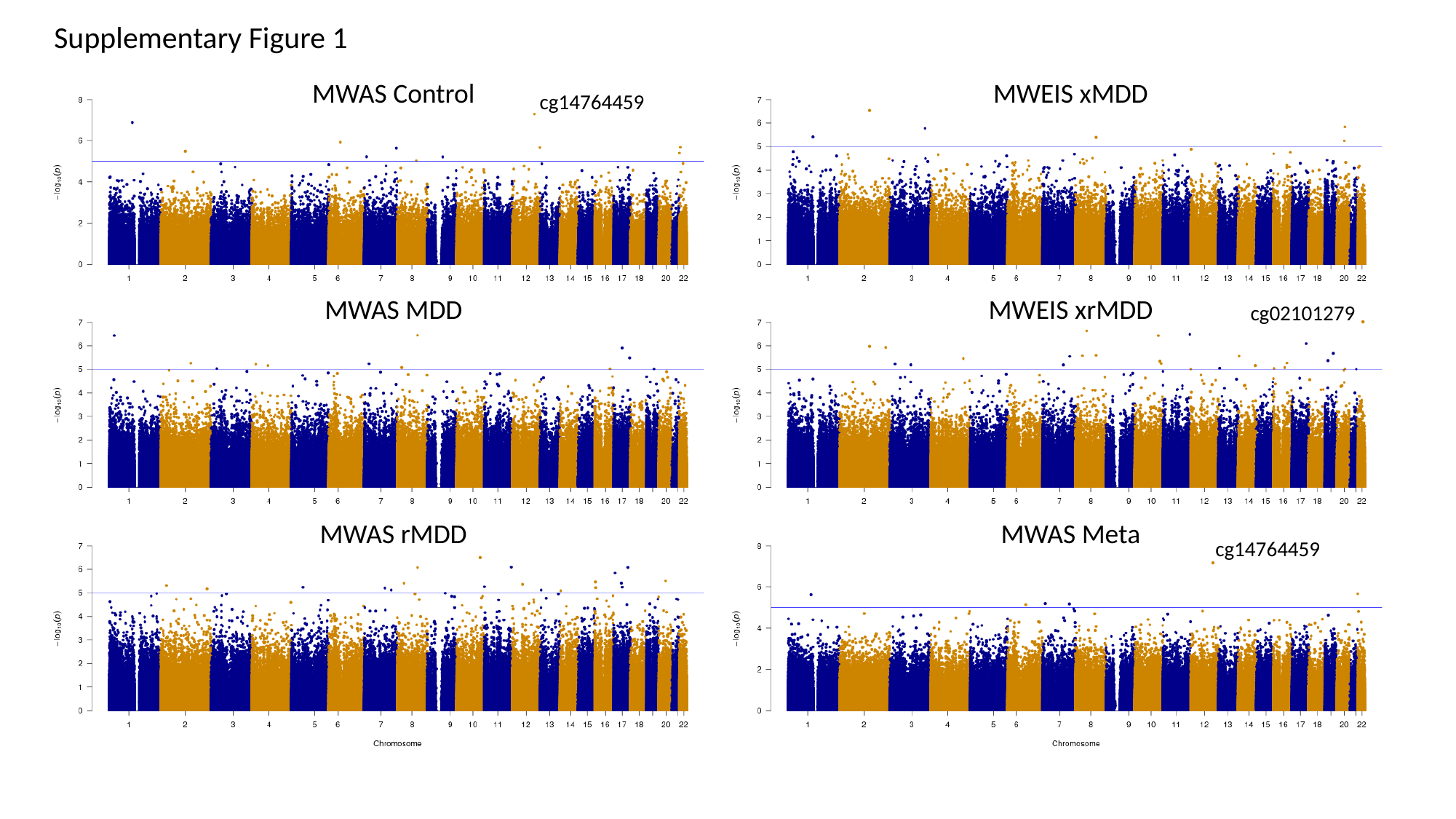

Supplementary Figure 1
MWAS Control
MWEIS xMDD
MWAS MDD
MWEIS xrMDD
MWAS rMDD
MWAS Meta
cg14764459
cg02101279
cg14764459

### Slide 2
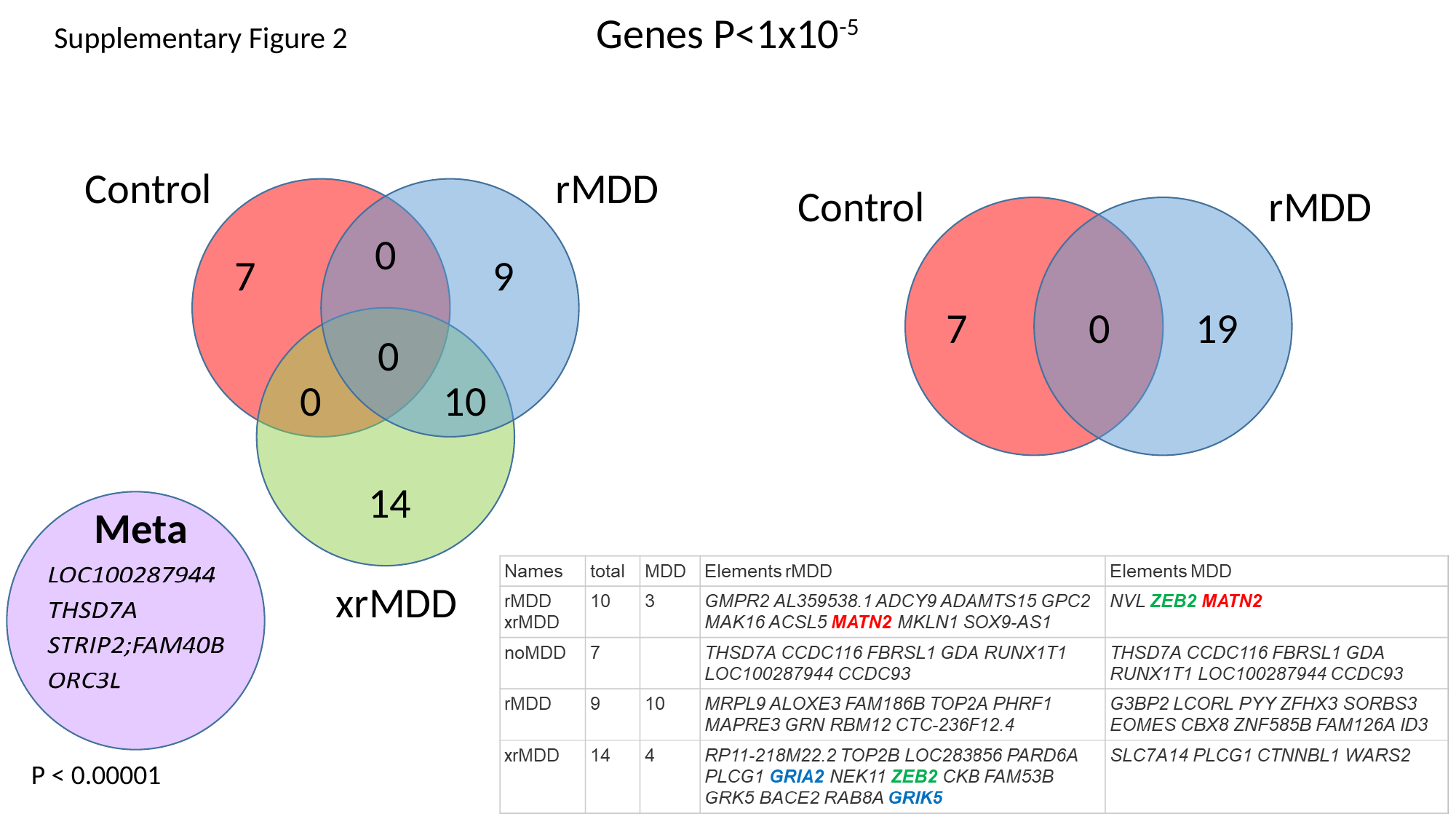

Genes P<1x10^-5^
Supplementary Figure 2
Control
rMDD
0
7
9
0
0
10
14
xrMDD
Control
rMDD
7
0
19
Meta
P < 0.00001

### Slide 3
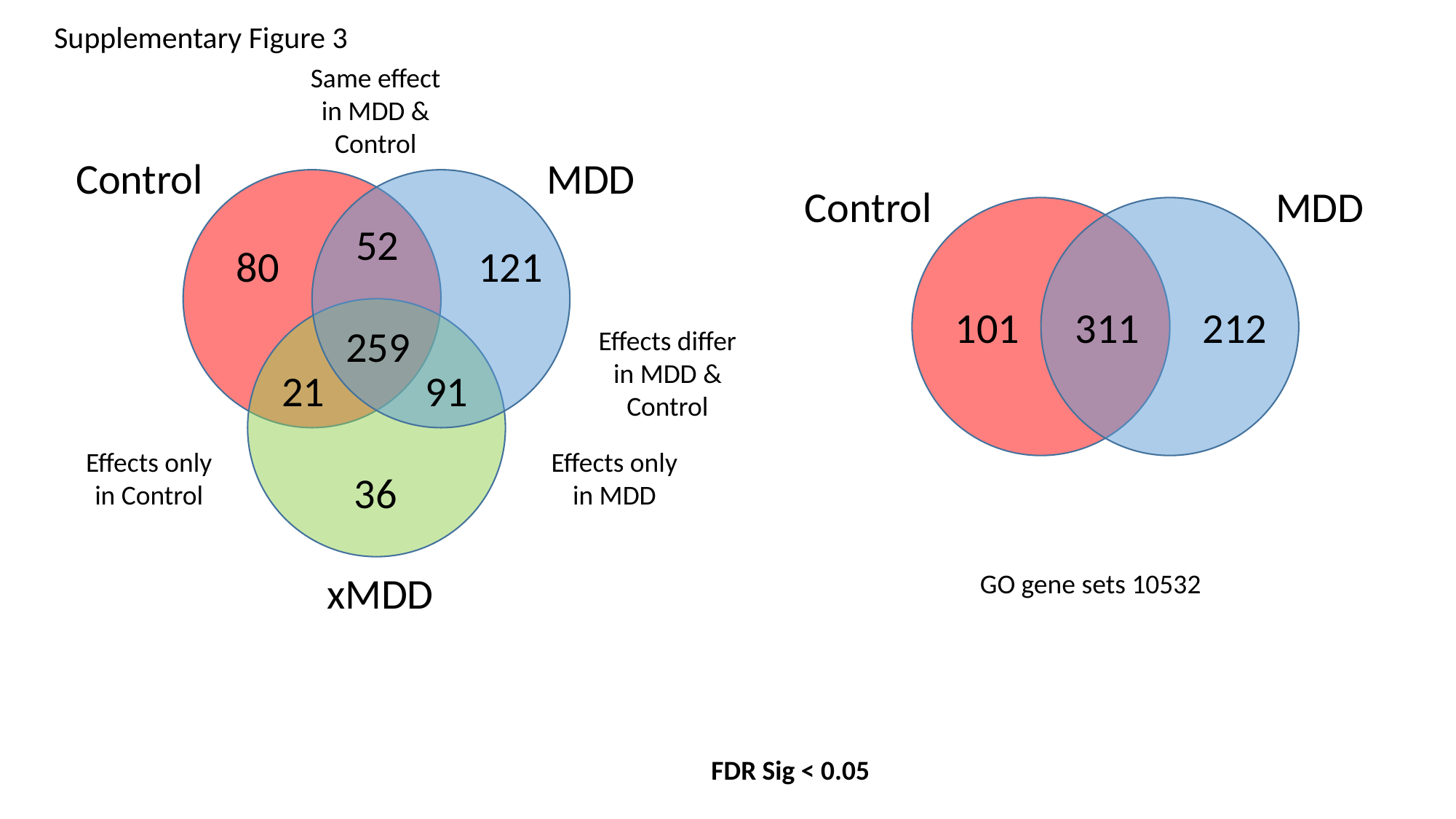

Supplementary Figure 3
Same effect in MDD & Control
Control
MDD
52
80
121
259
21
91
36
xMDD
Control
MDD
101
311
212
Effects differ in MDD & Control
Effects only in Control
Effects only in MDD
GO gene sets 10532
FDR Sig < 0.05

### Slide 4
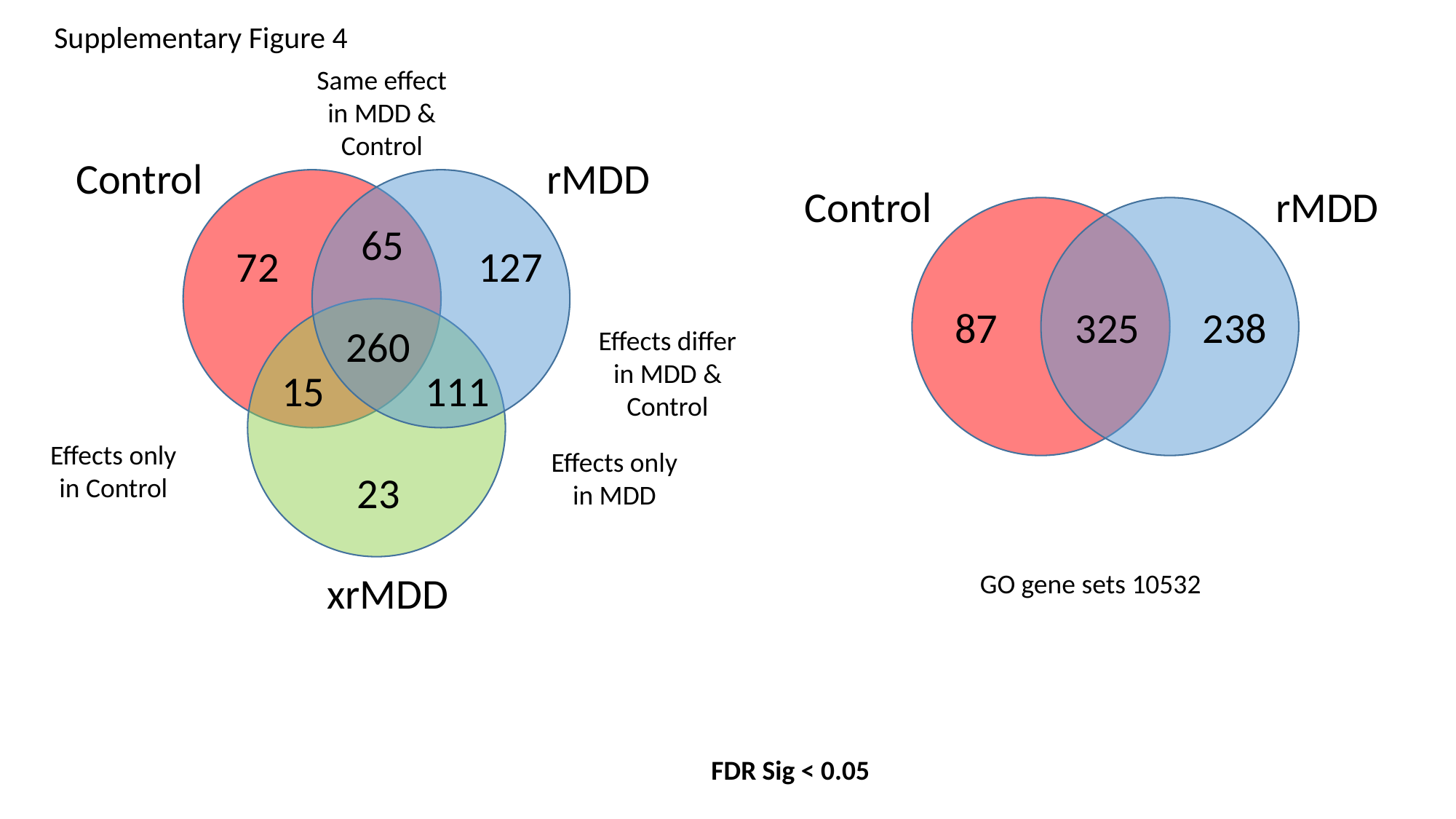

Supplementary Figure 4
Same effect in MDD & Control
Control
rMDD
65
72
127
260
15
111
23
xrMDD
Control
rMDD
87
325
238
Effects differ in MDD & Control
Effects only in Control
Effects only in MDD
GO gene sets 10532
FDR Sig < 0.05

### Slide 5
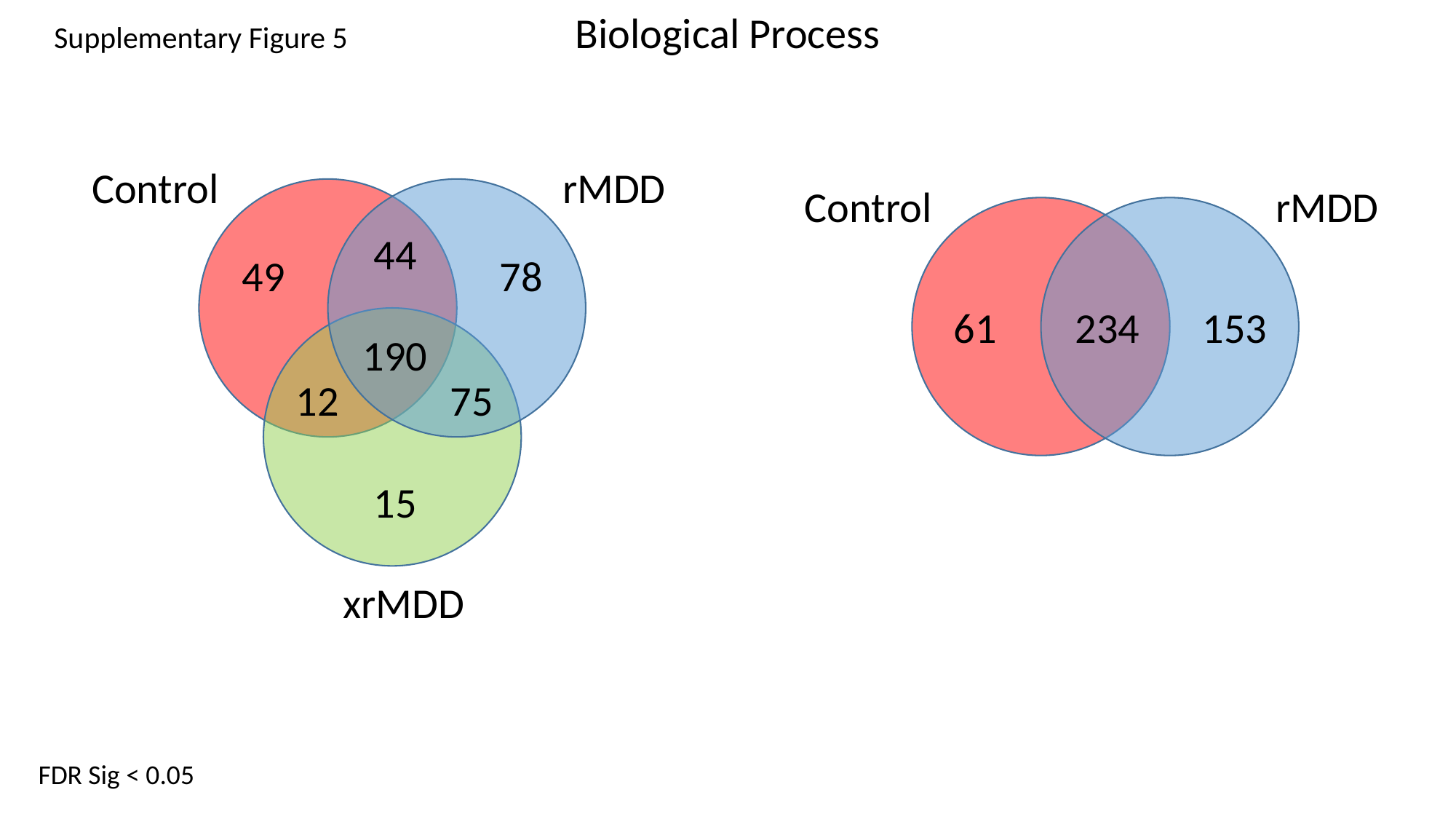

Biological Process
Control
rMDD
44
49
78
190
12
75
15
xrMDD
Supplementary Figure 5
Control
rMDD
61
234
153
FDR Sig < 0.05

### Slide 6
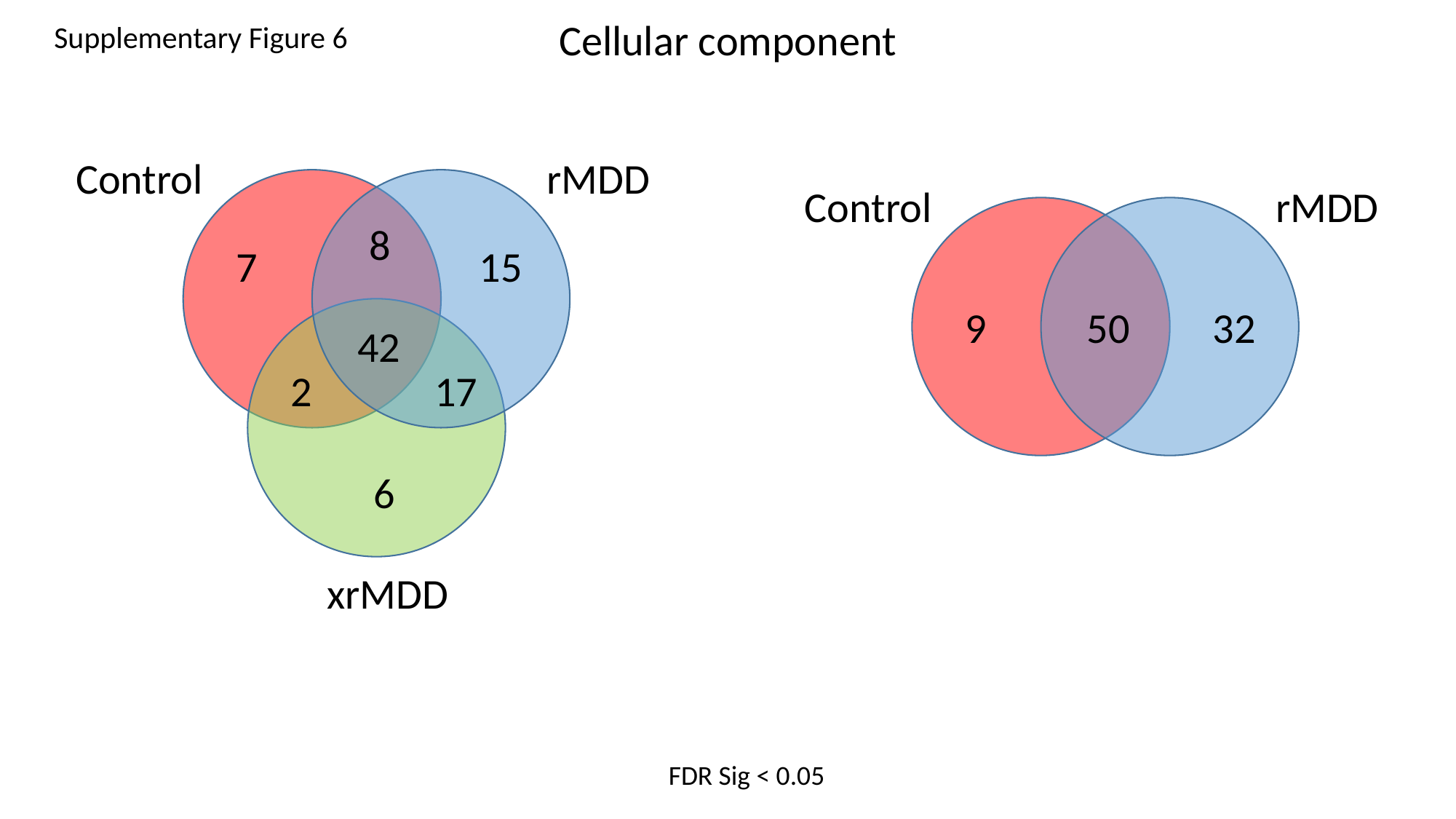

Cellular component
Control
rMDD
8
7
15
42
2
17
6
xrMDD
Supplementary Figure 6
Control
rMDD
9
50
32
FDR Sig < 0.05

### Slide 7
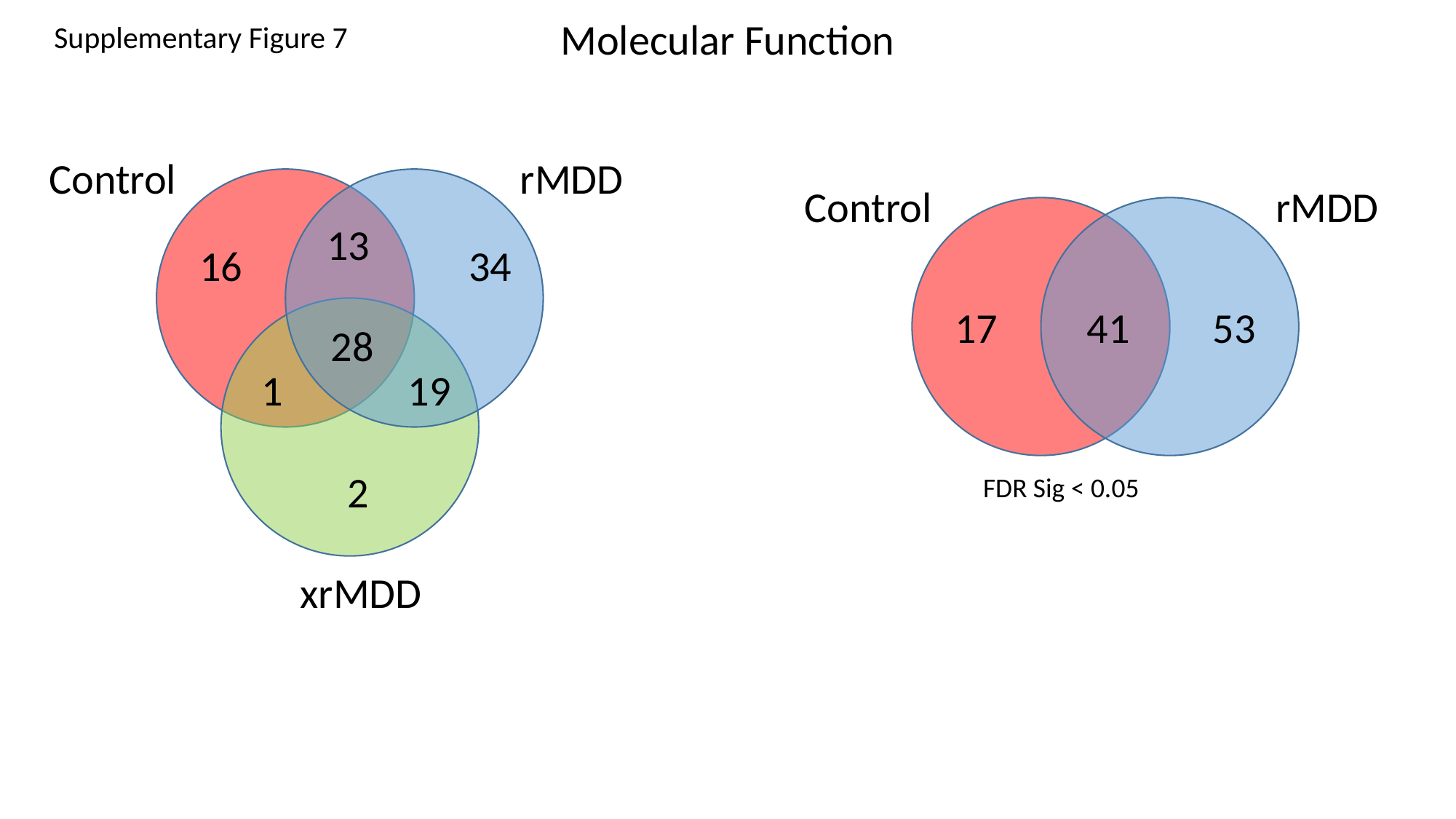

Molecular Function
Control
rMDD
13
16
34
28
1
19
2
xrMDD
Supplementary Figure 7
Control
rMDD
17
41
53
FDR Sig < 0.05

### Slide 8
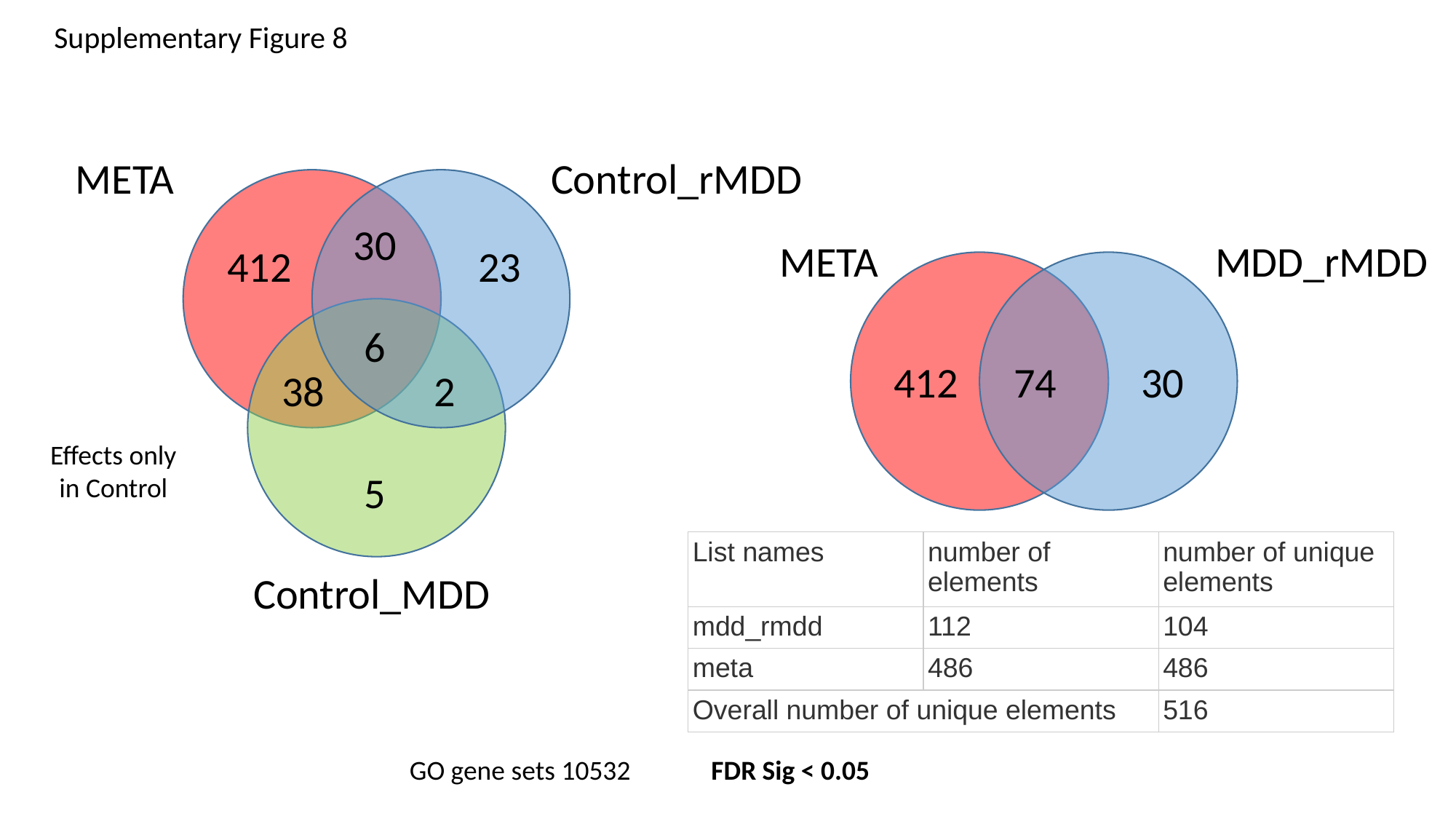

Supplementary Figure 8
META
Control_rMDD
30
412
23
6
38
2
5
Control_MDD
META
MDD_rMDD
412
74
30
Effects only in Control
| List names | number of elements | number of unique elements |
| --- | --- | --- |
| mdd\_rmdd | 112 | 104 |
| meta | 486 | 486 |
| Overall number of unique elements | | 516 |
GO gene sets 10532
FDR Sig < 0.05

### Slide 9
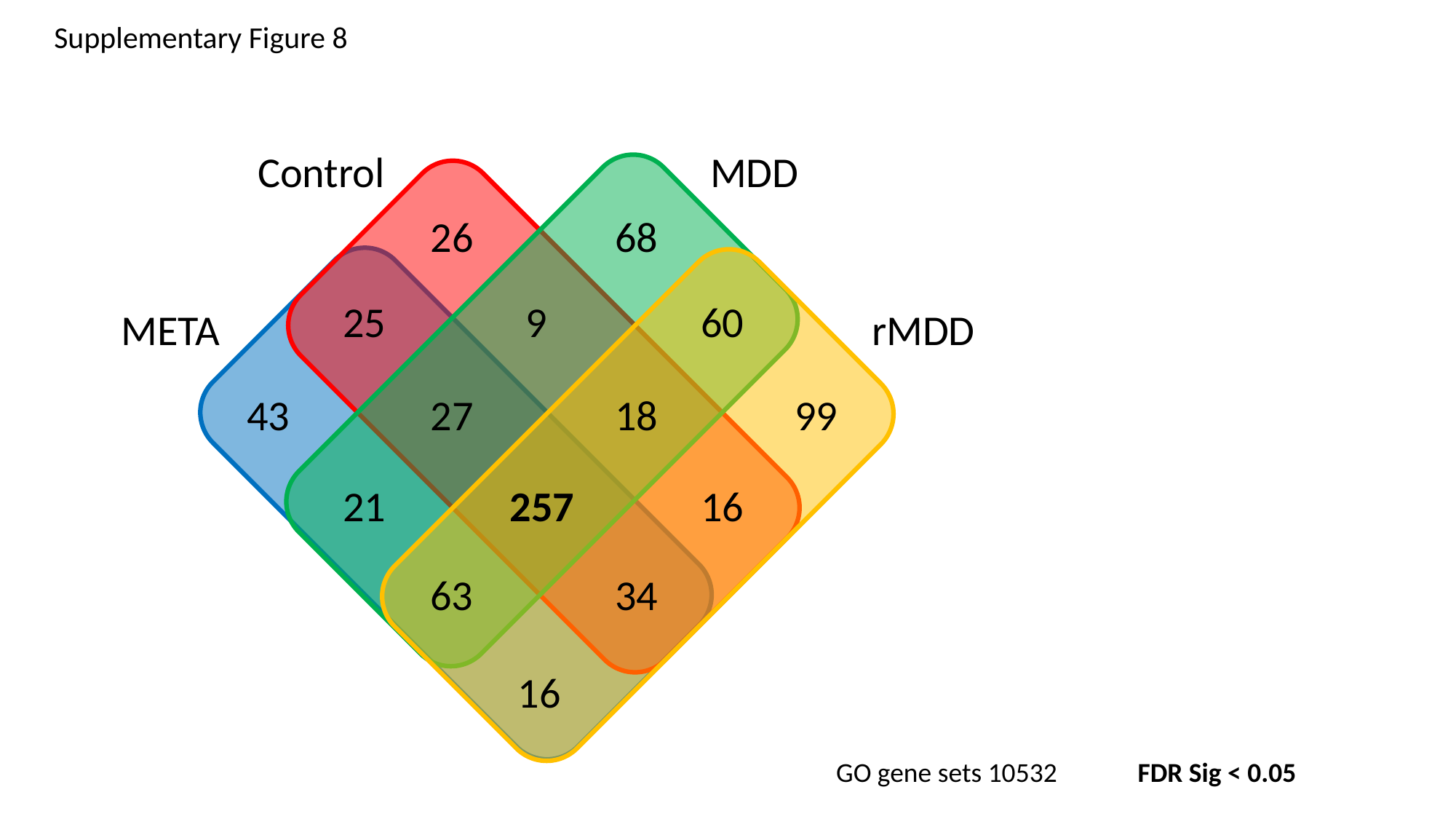

Supplementary Figure 8
Control
MDD
26
68
25
9
60
META
rMDD
43
27
18
99
21
257
16
63
34
16
GO gene sets 10532
FDR Sig < 0.05

### Slide 10
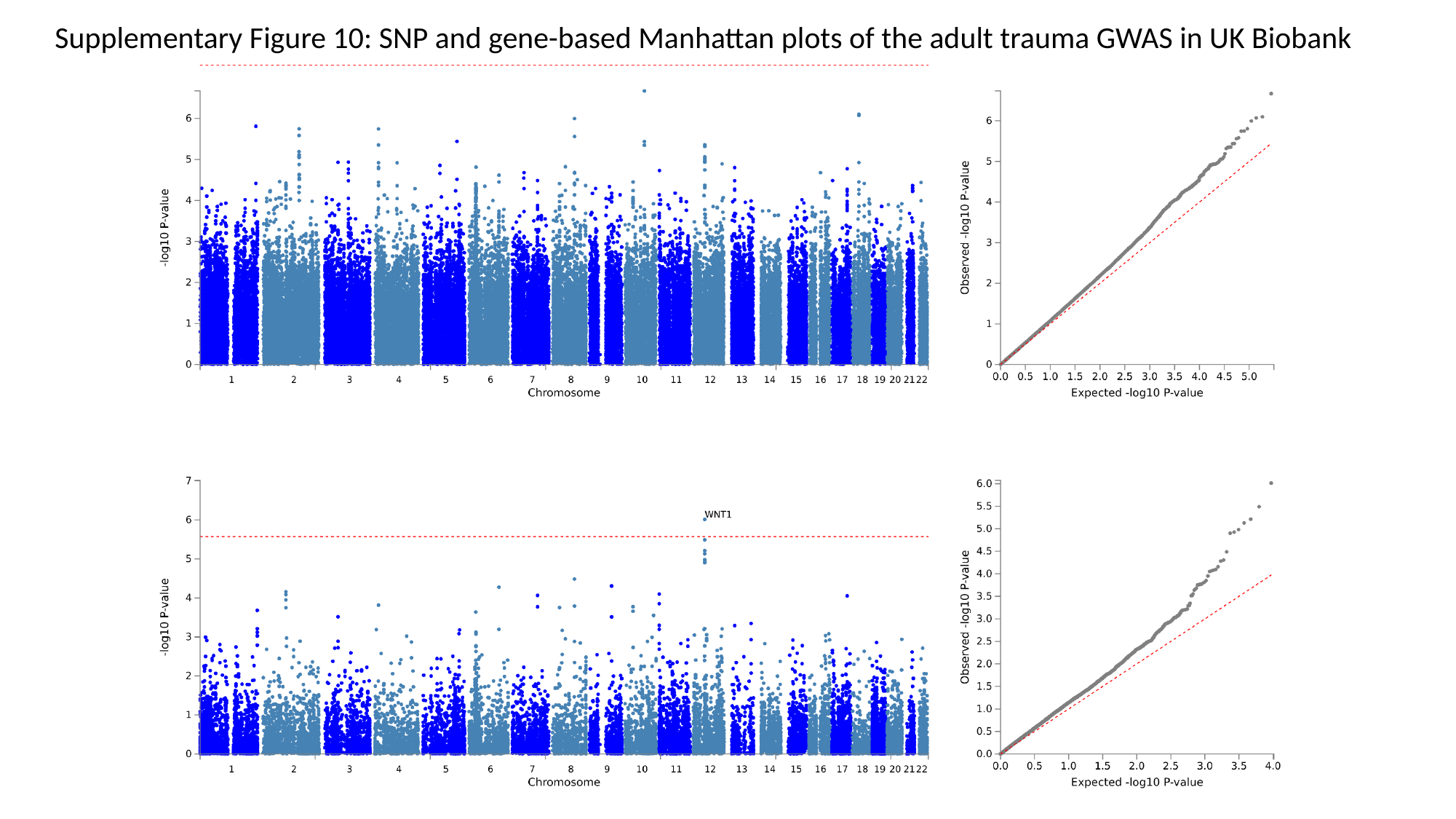

Supplementary Figure 10: SNP and gene-based Manhattan plots of the adult trauma GWAS in UK Biobank

### Slide 11
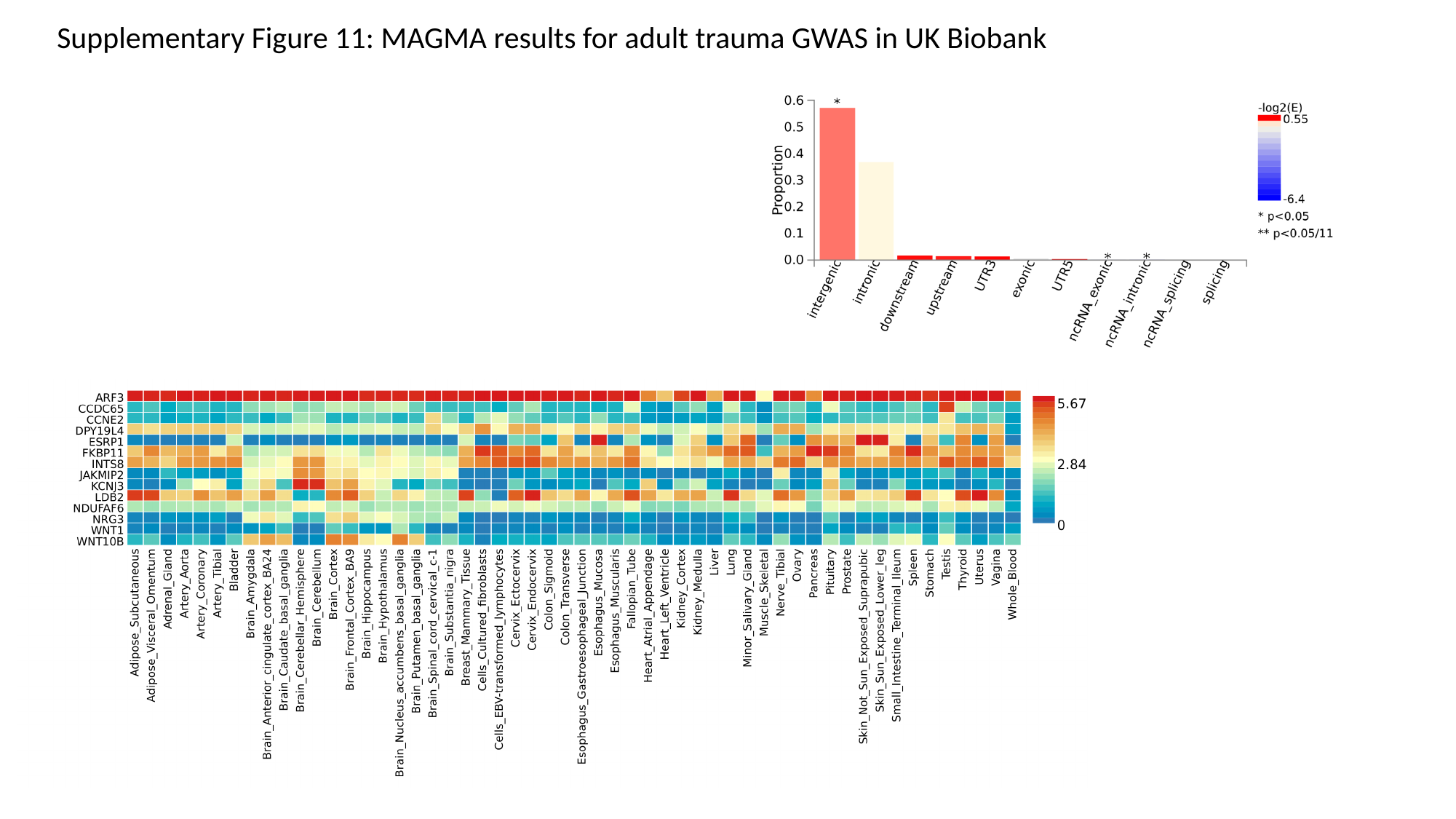

Supplementary Figure 11: MAGMA results for adult trauma GWAS in UK Biobank

### Slide 12
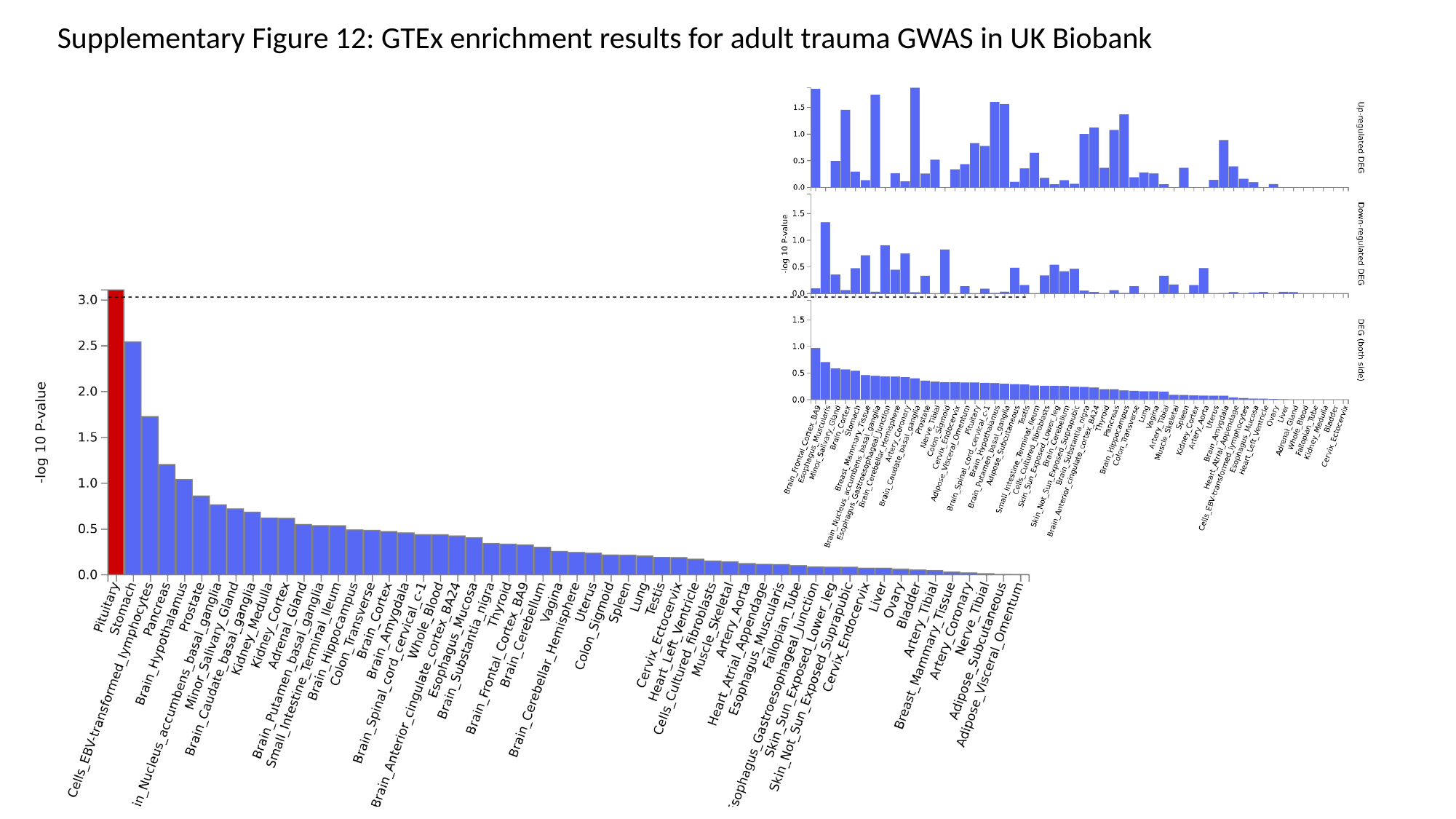

Supplementary Figure 12: GTEx enrichment results for adult trauma GWAS in UK Biobank
